# Inter individual variability in neuronal expression of heat shock protein genes predicts stress survival in *Caenorhabditis elegans*

**DOI:** 10.1101/2023.06.28.546835

**Authors:** Pia Todtenhaupt, Sharlene Murdoch, Catalina A. Vallejos, Olivia Casanueva, Laetitia Chauve

## Abstract

Despite being isogenic and grown under controlled conditions, *C. elegans* populations exhibit widespread inter-individual variability in many traits, making it an ideal model organism to investigate non-genetic influences on phenotypic diversity. Our particular interest is to study the consequences of inter-individual variability in genes encoding heat shock proteins, which are expressed at low levels under non-stimulated conditions. To robustly quantify inter-individual gene expression, we developed a novel pipeline that combines a highly efficient cDNA extraction method with a high-throughput qPCR nanofluidics technology with a bespoke computational analysis. We validated our approach by benchmarking against *in vivo* reporters. We also screened among hundreds of stress inducible genes, and identified a regulon formed by transcripts belonging to the inducible heat shock protein family. We demonstrate, using a bipartite *in vivo* fluorescent reporter, that the inter-individual variability in the stress regulon stems mostly from anterior neurons. Our studies demonstrate for the first time that, under physiological and unstimulated conditions, the variable expression of neural stress responses has cross-tissue consequences for fitness at the individual worm level, suggesting an adaptive role under variable environmental conditions.

## INTRODUCTION

Having the same genotype, and being exposed to the same environmental conditions, does not guarantee a shared, unique phenotype. Stochastic and micro-environmental differences in gene and protein expression occur in bacterial to human cells (Hardo and Bakshi 2021; Bashkeel et al. 2019) and in the multicellular model organism, *C. elegans*, which is considered to be isogenic (Casanueva, Burga, and Lehner 2012; Mendenhall et al. 2012; Cypser et al. 2013). Gene expression variability is pervasive even in isogenic worms, calling for multiple buffering mechanisms during critical stages of development (Huelsz-Prince et al. 2022; Traets et al. 2021) and driving lifespan differences among individuals (Kinser et al. 2021). In unicellular organisms, variability in gene expression enhances fitness in a fluctuating environment, as an adaptive strategy that anticipates environmental change and enhances survival (Blake et al., n.d.; Acar, Mettetal, and Van Oudenaarden 2008; Kinser et al. 2021).

Another example of variability in gene expression includes stress-induced expression of heat shock proteins *(hsps),* also called molecular chaperones. Heat shock proteins (*hsps*) are a family of cytosolic proteins, originally described as being produced by cells in response to heat stress (Ritossa 1962), but are they also induced by other stressors in all living organisms, from bacteria to humans (Lindquist 1986). Members of this group perform chaperoning functions by stabilizing new proteins and proteins damaged by stress. Their transcriptional induction is a key part of the heat shock response and is induced primarily by the transcription factor, Heat shock factor 1 (*hsf-1*) (Morimoto 1998). Although HSF-1 acts cell autonomously to protect each cell of an organism, the overexpression of HSF-1 in neurons can potentiate an organism’s responses to exogenous stress in peripheral tissues (Douglas et al. 2015). A key limitation of some of the genetic manipulation approaches is that over-expression tools can push the system to a possible, yet unlikely, physiological state. The range of possible physiological solutions that organisms deploy *in vivo* to coordinate a stress response across tissues remains currently unknown.

Despite being essential to cells and showing a high degree of dose sensitivity, *hsp* expression is known to vary among individuals in a population. In the case of yeast cells, *hsps* respond to the variability of their common upstream regulator *Mns-2/4*. In fact, steady state *hsp* levels correlate with a yeast cell’s response to an exogenous stimulus and more importantly, predict the survival of individual yeast cells to exogenous stressors (Stewart-Ornstein, Weissman, and El-Samad 2012). In *C. elegans*, an induced stress response has variable, dose-dependent outcomes. For example, worms with higher, heat-induced HSP protein levels have a more robust response to environmental and genetic perturbations, at the cost of fecundity (Casanueva, Burga, and Lehner 2012). Such dose-dependent consequences of higher *hsp* levels in individual worms highlight the trade-off underlying *hsp* expression levels and the potential adaptive value of variable *hsp* expression at the population level. Less is known, however, about the extent of *hsp* expression variability under steady state, unstimulated conditions, and the consequences of this variability for fitness. This is particularly challenging for highly inducible *hsps*, as their basal expression is very low.

Our goal was to develop a method to reliably measure inter-individual variability in gene expression in isogenic *C. elegans* designed specially to monitor variability for lowly expressed genes, such as *hsps*. However, a main challenge of single-worm (or single cell) transcript detection technology is that when starting from small amounts of material, the capture efficiency is lower compared to bulk samples. This is problematic for rare transcripts for which technical noise is more prominent and that may fall below experimental limits of detection. Therefore, the reliable detection of rare transcripts, such as highly inducible *hsps*, requires the optimization of methods with higher sensitivity and reproducibility. To do so, we have previously developed a highly sensitive, multiplexed nanofluidics qPCR method that provides both high experimental throughput and a wider dynamic range of detection (Chauve et al. 2020). We also use a Bayesian statistical model to infer transcriptional variability estimates that take into account technical censoring and spurious technical noise.

We validate our inter-individual variability measurements using transcriptional reporters, which provide a gold standard of stable and highly variable genes. We apply this pipeline to screen expression of hundreds of stress response genes, we identify a regulon of low expressed inducible *hsps* that are highly variable in the absence of overt stress. To investigate the physiological consequences of variability in inducible *hsps*, we then use a binary expression system to follow the activity of the variable *hsp-16.41* promoter and single molecule RNA fluorescent in situ hybridization (smRNA-FISH). We discovered that variability in inducible *hsp* expression in *C. elegans* stems primarily from their neurons. The number of stressed neurons predicts survival to stress, in line with observations that have causally linked *hsf-1* expression in neurons with the control of peripheral stress responses (Douglas et al. 2015). These results thus indicate that neuronal stress not only can induce a survival response but that it does so under physiological conditions.

## RESULTS

### An improved pipeline to measure inter-individual variability from single-worm PCR gene expression data applied to stress response genes

Our aim was to quantify transcriptional variability for genes known to regulate proteostasis under unperturbed conditions. We screened 168 stress related genes (**Table S1**) under unstimulated conditions. The mRNA levels of inducible *hsps*, such as *hsp-16.2, hsp-16.41 and hsp-70(C12C8.1)*, range from 2 to 58 attomol/µL, levels that are undetectable using standard methodologies. To reliably detect low quantities of RNA, we recently optimized a fast and robust method to prepare cDNA from single worms, called “Worm-to-Ct” (Chauve et al. 2020). The method’s increased accuracy and sensitivity relies on the omission of the isolation step of easily degradable RNA during cDNA synthesis and on the introduction of a pre-amplification step. Using Worm-to-Ct protocol, we are able to capture between 2 to 4 fold more cDNA from single worms as compared to trizol extraction on single worms (**Figure S1A-B**). We coupled “Worm-to-Ct”, with multiplexed, nanofluidics qPCR technology to increase the experimental throughput and to provide a broader dynamic range of detection, starting from input concentrations of ∼2 attomoles/mL RNA (Chauve et al. 2020). **Figure 1A** recapitulates the Worm-to-Ct method and further technical details can be found in the supplemental methods section.

**Figure 1:**
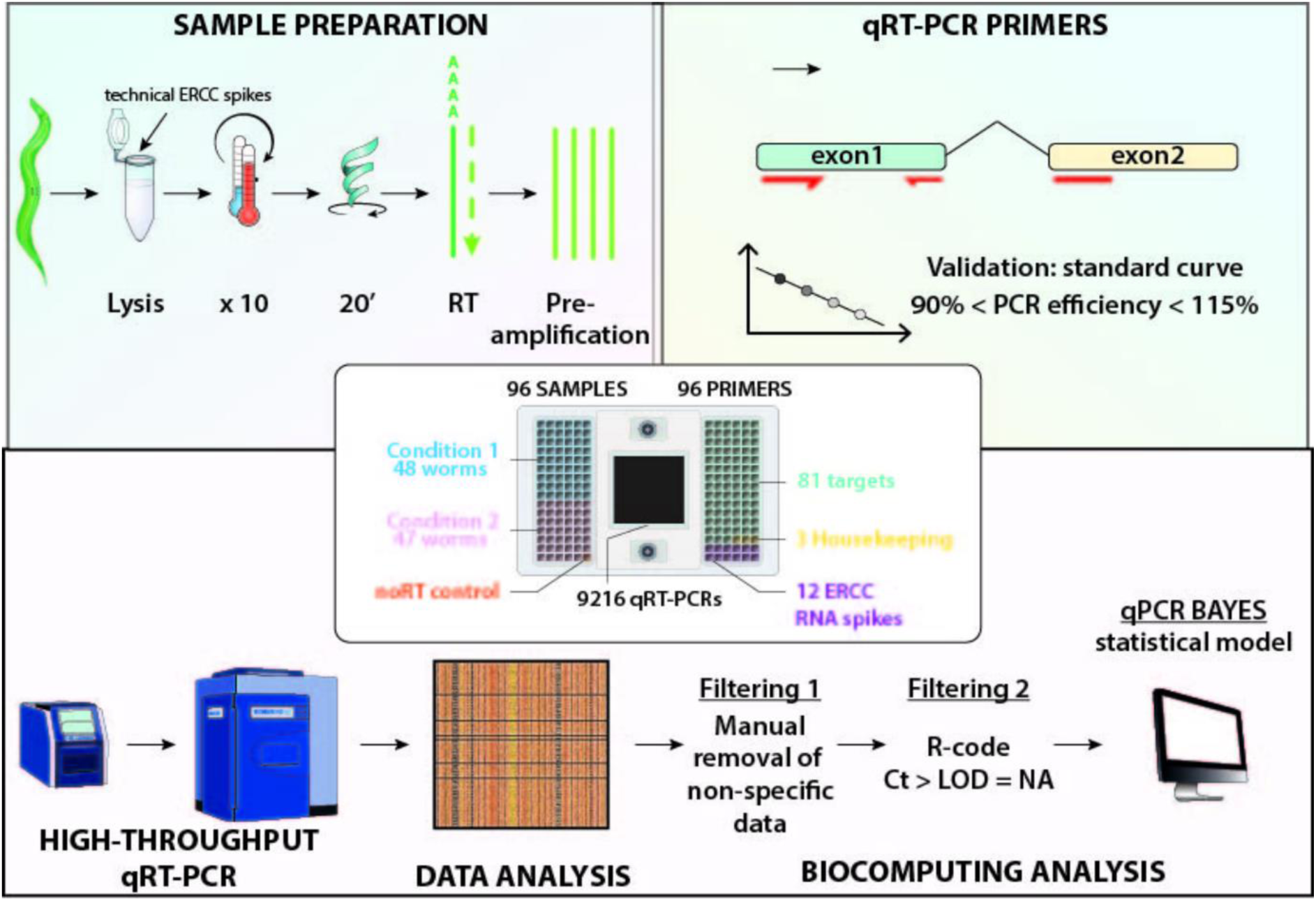
Overview of the method and key controls used to quantify mRNA transcripts from single worms. Overview of the pipeline used in this study to estimate gene expression variability using the Worm-to-Ct qPCR method coupled to qPCR Bayes statistical model. The single worm cDNA preparation called “Worm-to-Ct” protocol is described in details in (Chauve et al. 2020). List of genes amplified is provided in Table S6. The main steps of the “Worm-to-Ct” protocol added to the specificities proper to measure inter-individual variability (e.g. adding synthetic RNA-spike in, filtering data pst-run) are detailed in the general method section of this paper, under sample preparation. The dynamic range of detection of this method is higher than that obtained by using standard qPCR methods and comparable to that of other nanofluidic-based assays (Devonshire, Elaswarapu, and Foy 2011; Chauve et al. 2020). The experimental reproducibility is achieved by the consistent minimization of technical sources of variability, including the use of a pre-amplification step, eliminating the need for technical triplicates (Chauve et al. 2020). In this study, we performed essential controls to test the sensitivity (limit of detection, Figure S1C-D), the specificity (absence of genomic DNA contamination, Figure S1F-I) of the Worm-to-Ct method, and to ensure our qPCR methods follows previously determined PCR guidelines (Bustin et al. 2009). Furthermore, the primers used in this study (listed in Table S2) were tested for both specificity and efficiency (see general methods). We used the geNorm algorithm (Vandesompele et al. 2002) to determine cdc-42, pmp-3 and ire-1 as the most stable housekeeping (HK) (see supplemental methods and table 1). We show in figure S1A-B, that the Worm-to-Ct method surpasses traditional Trizol extraction methods by increasing the capture efficiency of transcripts from single-worms. We determined a reasonable number of worms (n=45) to accurately monitor inter-individual variability in gene expression (Figure S2).

**Table 1:** Single worm gene expression data.

| Figure | plate | condition | stage | Strain and genotype |
| --- | --- | --- | --- | --- |
| Fig 2A,2D | plate 23 | sample 2 | day 1 adult | BCN1050 *crgIs1002[daf-21p::mcherry]* |
| Fig 2D | plate 20 | sample 2 | day 1 adult | MOC108 *bicSi3[daf-21p::daf-21]* |
| Fig 3A | plate 9 | sample 1 | day 1 adult | MOC1 *mgIs49[mlt-10p::GFP-pest] x7* |
| Fig 3B, 3C, 3D | plate 9 | sample 1-2 | day1/day2 | MOC1 *mgIs49[mlt-10p::GFP-pest] x7* |
| Fig 4G | plate 24 | sample 1 | day 1 adult | PS7171 *hsp-16.41p::cGAL/UAS::GFP* |
| Fig 4H | plate 24 | sample 2 | day1 adult | PS7167 *unc-47p::cGAL/UAS::GFP* |
| Fig S5Aleft-B | plate 11 | sample 1-2 | day1/day2 | MOC1 *mgIs49[mlt-10p::GFP-pest] x7* |
| Fig S5Amiddle-C | plate 12 | sample 1-2 | day1/day2 | MOC81 *mgIs49; gon-2(q388)* |
| Fig S5Aright-D | plate 14 | sample 1-2 | day1/day2 | MOC2 *mgIs49; glp-4(bn2)* |
| Fig S1 B | plate 15 | sample 1 | day 1 adult | MOC1 *mgIs49[mlt-10p::GFP-pest] x7* |
| Fig S3 A-B | plate 21 | sample 1 | day 1 adult | MOC1 - TRIzol extraction |
| Fig S3 A-B | plate 21 | sample 2 | day 1 adult | MOC1 - Worm-to-Ct protocol |
| Fig S3 C,D,E | plate 23 | sample 2 | day1 adult | BCN1050 *crgIs1002[daf-21p::mcherry]* |
| Fig S2 | plate 7 | 1 sample | day1 adult | MOC1 - 1 condition on the chip (90 worms) |

In qRT-PCR assays, gene expression is measured by *threshold cycles* (Ct), defined as the number of PCR amplification cycles that were required to achieve a given fluorescence level. Technical censoring arises when Ct values exceed a pre-specified limit of detection (typically set to 22 cycles) above which measurements are deemed to be unreliable or where the fluorescence threshold was not achieved (**Figure S1B-C**). Existing qRT-PCR analyses sequentially apply two pre-processing steps prior to downstream analyses. Firstly, censored values are imputed (typically using the experimental limit of detection, e.g. 22 cycles) and subsequently, the data is normalized to remove unwanted sample-specific effects. Imputation and the presence of technical noise can substantially affect inter-individual gene expression variability estimates. To address this, we have adapted a linear modelling framework to explicitly account for censoring, whilst using control genes to perform normalization and quantify technical noise (see Supplementary Methods).

### Validation of inter-individual variability measurements *in vivo*

To benchmark the estimates of inter-individual variability obtained from our pipeline (**Figure 2A**), we compared them to in *vivo* transcriptional reporters that faithfully represent the transcriptional output of endogenous genes (**Figure 2B-C, Figure S4 A-D**). The variability of these “gold standards” (GS) was captured by the coefficient of variation (CV), defined as the standard deviation divided by the mean, calculated from fluorescence intensity measurements of transcriptional reporters. The CV values of these endogenous genes revealed that they were either highly stable across individual worms (*hsp-1* and *unc-54* with CV values of 0.14 and 0.1 respectively), or highly variable across individual worms (*ttr-45* and *nlp-29*, with CV values of 0.39 and 0.83 respectively) (**Figure 2C, Table S4**). Furthermore, inter-individual variability estimates were positively correlated with the CV of gold standards (Pearson correlation coefficient R^2^=0.92), indicating a high degree of concordance between the two methods (**Figure 2D**). In subsequent figures, a threshold was set such that inter-individual variability estimates were considered high when values surpassed *ttr-45* inter-individual variability estimate (i.e. CV > 0.39).

**Figure 2.**
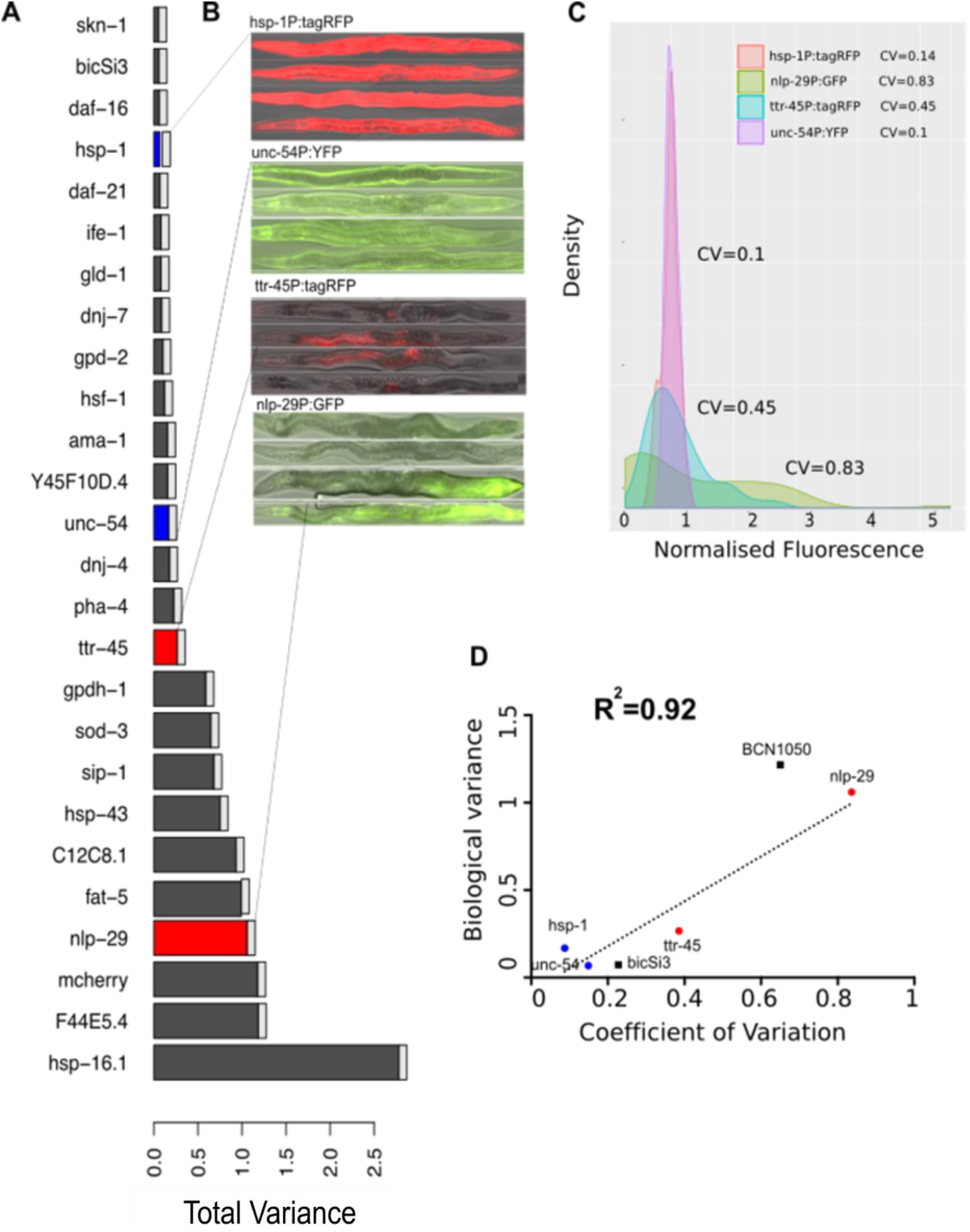
Benchmarking inter-individual variability estimates derived from our pipeline. **(A)** Comparison between inter-individual variability estimates obtained from our pipeline, applied to day 1 young adults (raw data: plate 20_2, tables S2 and S3) and **(B-C)** Coefficient of variation (CV) values – defined as the standard deviation divided by the mean and reported in Table S4 – were obtained from the following transcriptional reporters: MOC92 *bicIs10[hsp-1p::RFPAI]*, MOC119 *bicls12[ttr-45p::tagRFP]*, IG274 *frIs7[nlp-29p::GFP + col-12p::dsRed]* (Pujol et al. 2008) and AM134 *rmIs126[unc-54p::YFP]* (Morley et al. 2002). Animals were imaged at the young adult stage on an epifluorescence microscope. Day 2 adult animals were used in the case of IG274 and MOC119, as the transgene was not visible at young adult stage. To ensure transcriptional reporters depicted in B faithfully represent endogenous mRNA levels, qPCR was performed to ensure that the fluorophore mRNA driven by the transgene correlates to the levels of endogenous genes transcripts (Figure S4) (**D)** Correlation between coefficient of variation (CV) values obtained from fluorescence measurements and inter-individual variability obtained by our pipeline. (**A, D**): Red color indicate “variable gold standards” *nlp-29* and *ttr-45*, which are highly variable by both methods. Blue dots indicate “stable gold standards” *hsp-1* and *unc-54*, which are stable by both methods. In D, BCN1050 and bicSi3 represent additional variable and stable transcriptional reporters that were assessed. BCN1050 animals carry multicopy transgene *crgIs1002[daf-21p::mcherry]* while bicSi3 carry single copy transgene *bicSi3[daf-21p::daf-21].* Comparative analysis using fluorescence measurements (Figure S4, table S4) and inter-individual variability estimates obtained by our pipeline (Figure 2D) revealed that multicopy *daf-21* transcriptional reporters were highly variable while single-copy *daf-21* reporters were stable. Images, density distributions and CV of *daf-21* transcriptional reporters are given in figure S4 and table S4.

### Small heat shock protein expression is highly variable and hsps form an “inter-individual variability regulon” in the absence of exogenous stress

We estimated inter-individual variability in the expression of genes known to regulate proteostasis in young adult animals grown under standard unperturbed conditions and day 1 and day 2 of adulthood (**Figure 3A-B).** These genes were detected across a broad range of mean expression values (although higher inter-individual variability was generally found to be associated with lower mean Ct Values) and includes heat shock protein family members belonging to different classes (*hsp-60, hsp-100, hsp-40, hsp-90, hsp-70*), and transcription factors involved in the modulation of stress responses (**Figure 3A-C**). The gene group that passed the threshold for high inter-individual variability (set to be higher than that for *ttr-45*) was enriched for genes involved in inducible heat shock stress responses (*hsp-16* family and inducible *hsp-70*). We also included genes involved in innate immunity and responsive to fungi (Pujol et al. 2008) as positive controls, since a reporter strain of *nlp-29* has been previously shown to vary across worms (Pujol et al. 2008) and corresponds to one of our gold standards (**Figure 2**). As expected, the genes that encode immunity related peptides from a neuropeptide family (*nlp-29, nlp-31, nlp-34, nlp-27*), are highly variable and co-regulated (**Figure 3C-D**).

**Figure 3.**
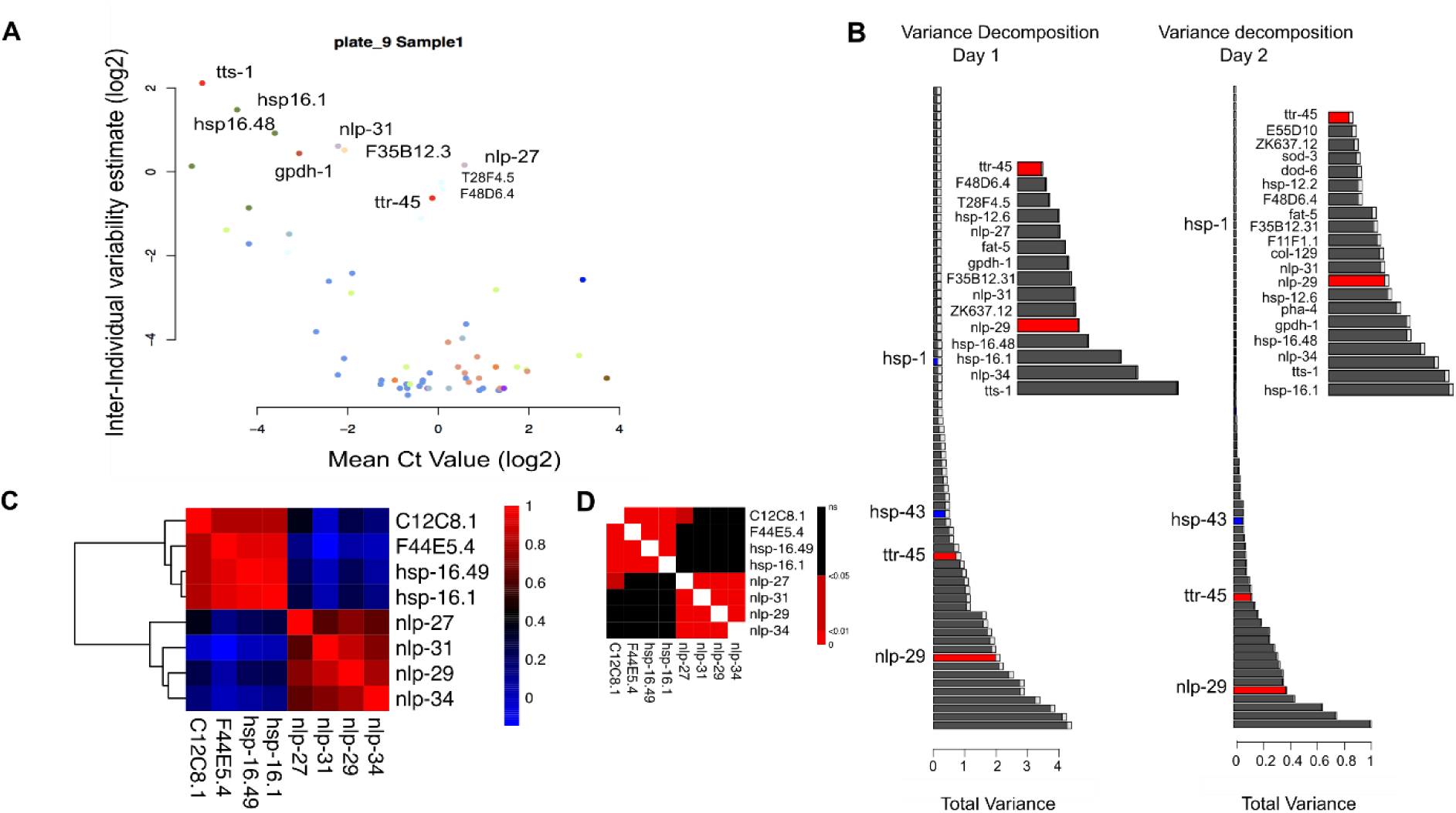
Stress response genes are highly variable from worm to worm in the absence of stress. **(A)** Scatter plot showing inter-individual variability of the Ct values with respect to the mean expression in wild type animals at day 1 of adulthood (plate 9_1, Table S2 and S3). An inverse relationship between mean expression and variability is evident. Note that Ct values on the X axis are plotted as 25 - Ct value, with 25 being an absolute maximal Ct value corresponding to no amplification on the Biomark (see Methods), so that lowest Ct values scale with mean expression. Each dot represents one target gene and the colors represent different families of related stress-response genes. **(B)** Variance decomposition at days 1 and 2 of adulthood. Gold standards are colored blue for stable expressed genes (*hsp-1*) and red for variable genes (*nlp-29* and *ttr-45*). Only genes that had higher variability estimates than *ttr-45* were considered as highly variable genes. Inter-individual variability is depicted in dark grey and technical variability as light gray, note that most of the measured variability is biological. Insets show the genes that scored high for inter-individual variability at the first two days of adulthood. They comprise highly genes from the small heat shock proteins family and genes encoding neuropeptides (*nlp-* family), involved in innate immunity. Note that overall inter-individual variability measured at day2 is lower than day 1 (the scale for inter-individual variability is different between the two days). **(C)** Clustered Pearson coefficient correlation matrix calculated on denoised Ct values obtained at day 1 of adulthood shown in (Plate 11_1, table S3). (**D**) Correlation Matrix depicting the corresponding Benjamini Hochberg (BH) adjusted p-values revealing significant correlation coefficients. Two distinct “transcriptional regulons”, with significant co-variation in gene expression can be observed for highly inducible heat response genes (*hsp-16.1, hsp-16.41*, *hsp-70* family genes C12C8.1 and F44E5.4), and innate immunity response neuropeptides (*nlp-27, nlp-29, nlp-31* and *nlp-34*). Genes within each transcriptional regulon correlate with p-values<0.01. Animals were synchronized at day 1 of adulthood, based on the fluorescent of the *mlt-10p::*GFP::PEST transgene (Frand, Russel, and Ruvkun n.d.), with transient peaks of fluorescence accompanying every molt (see general methods).

We then determined if steady state transcript expression levels in single worms reveal functional groups of genes that are known to respond in a coordinated fashion to common environmental cues and transcription factors, even in the absence of exogenous stressors. This sort of variability is normally referred to as *Pathway-related* variability (Stewart-Ornstein, Weissman, and El-Samad 2012). **Figure 3C-D** shows the Pearson correlation and adjusted p-value to determine statistical significance, which highlights a highly positive and significant correlation between members of each sub-cluster (correlation by p<0.01). Each group is known to respond to a specific transcription factor (TF) and to contain TF binding sites within their promoter regions. *Hsps* are collectively regulated by the transcription factor HSF-1 (Morimoto 1998) and the positive controls encoded by *nlps* genes are collectively regulated by GATA factors and by MAP kinase signaling (Pujol et al. 2008) and as expected, show a signature of co-regulation. We thus conclude from these results the coherence in gene expression at basal levels among individual worms corresponds to *pathway-related* inter individual variability.

### Inter-individual variability of *hsp* family stems from anterior neuronal cells

The standard heat shock-inducible transcriptional reporters do not produce detectable levels of fluorophores under basal conditions (Golden et al. 2020), precluding any further investigation on the potential adaptive significance of inter-individual variability. To circumvent this technical hurdle, we sought to amplify the expression of GFP driven by a heat shock promoter, by means of a binary expression system (Wang et al. 2017). **Figure 4A** shows GFP expression in animals containing two multicopy integrated transgenes: the first uses the promoter of *hsp-16.41* to drive cGAL (Gal4 adapted for expression in *C. elegans*), and the second contains a promoter with multiple cGAL binding sequences (upstream activated sequences, UAS), to drive GFP expression. When noticed that – contrary to the expectation of a rather ubiquitous somatic expression – *hsp16.41* activity was restricted to cells within the head region, (**Figure 4A**). If these GFP positive cells in the head are neurons, then they should co-express *rab-3*, a gene that is expressed pan neuronally (Stefanakis, Carrera, and Hobert 2015), (**Figure S6A-D**). As seen in **Figure S6,** GFP expression coming from most positive head cells co-localize with *rab-3p*::tagRFP expression, except for one cell which we identify as the head mesodermal cell (star in **Fig S6D**). Furthermore, the cells expressing GFP from *hsp-16.41p*::cGAL/UASp::GFP show axonal projections (**Figure S6C**, asterisks). This data suggests that at 20°C, the expression of *hsp-16.41* comes mainly from anterior neuronal cells.

**Figure 4:**
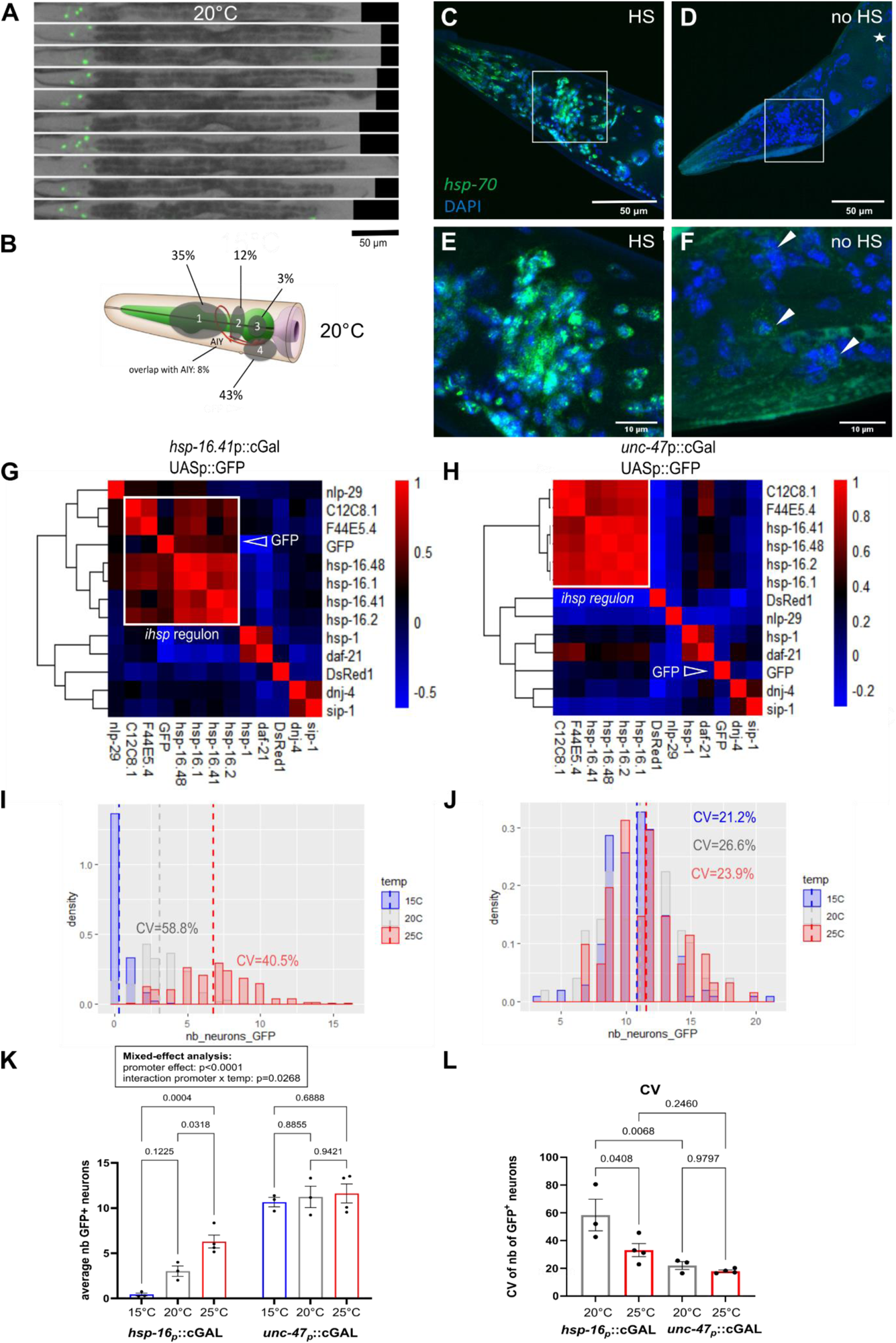
In the absence of overt stress, heat inducible hsps are expressed in neurons and their expression is highly variable and temperature sensitive. **(A)** A bipartite expression system (Wang, et al. 2017) (promoter::cGAL/UAS::GFP), was used to visualize the activity of the inducible heat shock promoter *hsp16.41* under unstressed conditions. Images taken at 5x magnification depicts PS7171 animals carrying both driver transgene *syIs400 [hsp16.41_p_::NLS::cGAL]* and effector transgene *syIs337 [15xUAS::pes-10::GFP]*. At 20°C, in the absence of heat stress, animals express GFP fluorescence only in cells within the head region of the worm, presumably neurons. **(B)** Diagram of the regions of the head where GFP expressing cells are present showing the percentage of GFP positive cells found in each area. The overlap with AIY neuron could be assessed as PS7171 animals also carry the co-injection marker *ttx-3_p_*::RFP, expressed in AIY. **(C-F)** Single molecule FISH (smFISH) was used to detect the presence of inducible *heat shock protein* transcripts in the presence (C, E) or absence (D,F) of heat shock using 63x confocal microscopy in day 2 animals. DAPI was used to highlight nuclei (blue); and the green channel shows the hybridization of a probe designed against an inducible *hsp-70* (C12C8.1). The pictures show a close-up of the head region of the worm. Whereas heat shock induces transcript production in all cells of the head (shown here in C, E) and of the body (Figure S5), in the absence of stress, *hsp-70* transcripts are primarily expressed in the head region of the worms (D, F and Figure S5). **(G-H)** Pearson correlation matrices for gene expression in PS7171 worms carrying *16.41_p_::cGAL/UAS_p_::GFP* **(G)** or PS7167 worms carrying the bipartite cGAL expression system *unc-47_p_::cGAL/UAS_p_::GFP* in GABAergic neurons **(H)**. GFP expression from *16.41_p_::cGAL/UAS_p_::GFP* activity correlates with the inducible *hsp* regulon in PS7171, while GFP expression from *unc-47_p_::cGAL/UAS_p_::GFP* activity does not correlate with inducible *hsps*. This indicates that the bipartite *16.41_p_::cGAL/UAS_p_::GFP* reporter faithfully correlates with endogenous inducible *hsps*. Dsred represents expression from the co-injection marker *unc-122_p_::DsRed1* present in both PS7171 and PS7167. Its expression does not correlate with *hsp* expression in both cases, as expected. **(I-J)** Histogram of distribution of the number of GFP expressing neurons in PS7171 animals carrying *16.41_p_::cGAL/UAS_p_::GFP* or PS7167 worms carrying *unc-47_p_::cGAL/UAS_p_::GFP* at different growth temperatures. Data is concatenated from three different biological replicates per condition (Table S5). Dash line represents mean expression. The transcriptional activity of *hsp-16.41* increases with temperature (I), while this is not the case for the *unc-47* expression in GABA-ergic neurons **(J)**. CV: coefficient of variation. The CV could not be calculated for *16.41_p_::cGAL/UAS_p_::GFP* expression at 15°C, as GFP expression is too low and the data is not normally distributed. **(K)** Mean number of GFP positive neurons in PS7171 and PS7167 animals raised at different temperatures. Mixed model analysis reveal that *hsp-16.41* and *unc-47* exhibit distinct effects (p<0.0001) with number of GFP expression neurons correlating with temperature in the case of *hsp-16.41* but not *unc-47* promoter. **(L)** Coefficient of variation of the number of GFP expressing neurons across biological replicates in PS7171 and PS7167 animals. One-way Anova.

The identity of these GFP-expressing neurons cannot be determined at the level of resolution used for this analysis. However, the location of GFP-positive cells varies from worm to worm, although they are most frequently found in the head regions highlighted in **Figure 4B**. In line with the high inter-individual variability, we detected for *hsp* transcripts, the number of GFP positive neurons varies highly from worm to worm (average CV= 0.59 at 20°C, **Figure 4I, L**), compared to the expression of GFP from a Gal4 system driven by *unc-47*, a promoter expressed in GABAergic neurons (average CV=0.26 at 20°C, **Figure 4J, L**). The observed data is consistent with a marked inter-individual variability in both the number and the location of neurons with an active stress response at 20°C. CV analysis performed from single-worm qPCR data further validated that the variability detected by the binary expression system faithfully reproduces the behavior of the entire *hsp* regulon when driven by *hsp-16.41* but not the *unc-47* promoter (**Figure 4G-H**).

The UAS-Gal4 system, adapted from yeasts, is known to be sensitive to growth temperatures (Wang et al. 2017). Although the cGAL system utilized in this work has been specifically adapted to robustly work at a range of temperatures in *C. elegans* (Wang et al. 2017), temperature may cause potential spurious effects on the cGal expression system. It has been established that *hsp16.41* responds to increments of temperature by the recruitment of HSF-1 to upstream promoter sequences (Douglas et al. 2015; Chauve et al. 2021), whereas *unc-47* promoter is not regulated by HSF-1. We hypothesize that the temperature dependent increase in the expression of *unc-47* should be negligible in comparison to that driven by *hsp16.41*. This hypothesis is correct when testing cGal driven by *hsp16.41* (**Figure 4I**, **K** and **Figure S6 F-G**) and not *unc-47* (**Figure 4J, K**). This shows that temperature sensitivity observed for *hsp-16.41* can be attributed to the nature of its promoter and is independent of the bipartite cGAL expression system. In terms of the effect of temperature on the variability of neuronal GFP driven by *hsp-16.41*, this is higher than neuronal GFP driven by *unc-47* at all temperatures tested (**Figure 4L**) and highest at 20°C (p=0.0408, **Figure 4L**). These experiments show that the expression driven by *hsp16.41* is higher and more reliable at warmer temperatures.

To further confirm the temperature sensitivity of *hsp* expression for endogenous transcripts we performed smRNA FISH using a probe against *hsp-16.41* mRNA at 15°C and 25°C. In this technique, endogenous *C. elegans* mRNA transcripts are labelled with a large number of complementary DNA oligonucleotides, and confocal fluorescence microscopy is then used to detect each mRNA molecule and its location inside the cell, as a diffraction limited spot (Femino et al. 1998; Raj et al. 2008). Members of the *hsp-16* gene transcripts are short (transcript length is 640 nucleotides for *hsp-16.41*). Therefore, when using a probe designed *hsp-16.41* it is also likely to target all members of the *hsp-16* gene family (see general methods). As expected, and consistent with results presented in **Figure 4**, we obtained *hsp-16* smRNA FISH signal in head cells, which was more pronounced at 25°C than at 15°C (**Figure S5H-L**).

Is the anterior neuronal expression of *hsps* specific to *hsp-16.41 or is it* generalizable to other endogenous inducible *hsps*? To answer this question, we used two methods: single-molecule RNA FISH (smRNA FISH) and spatial transcriptomics. For smRNA FISH, we used probes that hybridize to either an inducible *hsp-70 (ihsp-70)* transcript (C12C8.1, **Figure 5C-F**). As shown in **Figure 4C**, E most cells in the head, and in the body (**Figure S5, A, C, E,G),** express the *ihsp70* transcript after a pulse of heat shock. Consistent with the *in vivo* reporter studies, when the smRNA FISH is performed in the absence of heat shock, at 20°C, the primary cells that show positive hybridization to *ihsp70* (**Figure 5D, F** and **Figure S5 B, D,F,H)** are located in the head region. As an additional method to validate these observations, we looked at publicly available spatial transcriptomics data obtained from intact tissue sections (Ebbing et al. 2018). This spatial transcriptomic analysis showed that mRNA for *hsp* family members (*hsp-16.1 and hsp-16.11*) can only be detected from the head sections of hermaphrodite worms (**Figure S5 I-J).** Collectively the data indicates that under unperturbed conditions, the entire *hsp* regulon is activated in a variable number of anterior neuronal cells in single animals and sensitive to incremental changes in growth temperature.

**Figure 5.**
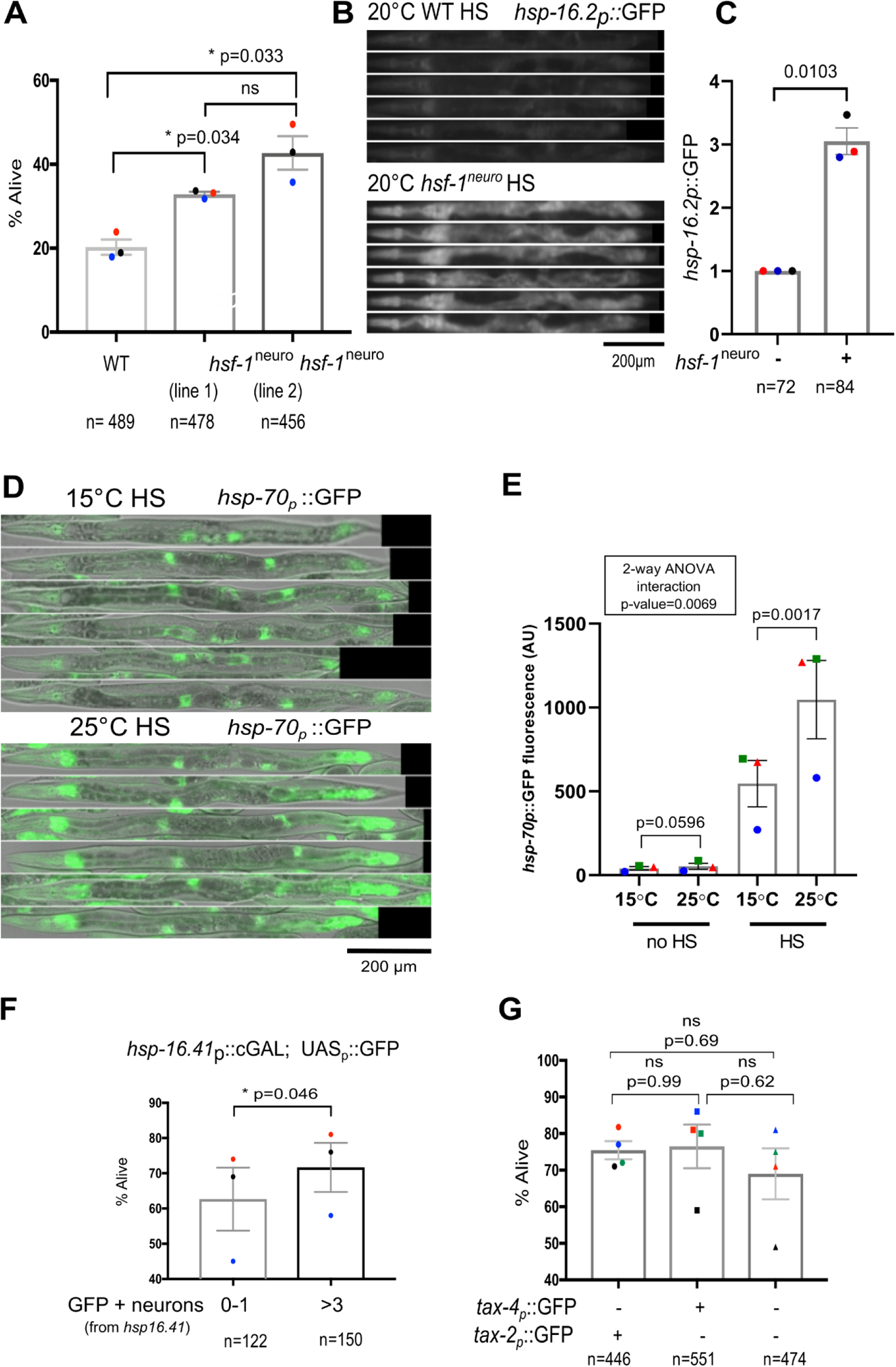
Neuronal expression of heat inducible transcripts can predict robustness to stress. **(A)** Two transgenic lines of animals overexpressing of *hsf-1* in neurons (*hsf-1^neuro^* lines1 and 2) exhibit increased thermotolerance with respect to wild type-animals as described (Douglas et al. 2015). Statistics was performed using a one-way Anova test. Thermotolerance assays were performed on young adult animals as described in material and methods. The Y-axis represents the percentage of worms surviving a three-hour heat shock. The points have been colour-coded by replicate pairs. **(B-C)** CL2070 and MOC153 animals carrying *dvIs70[hsp-16.2p::GFP]* and *uthIs368[rab-3p::hsf-1^neuro^]; dvIs70[hsp-16.2p::GFP]* respectively were heat shocked for 30 minutes at 34°C. The levels of GFP fluorescence were measured 3h post heat shock using a 5x objective inverted macro-zoom. Fluorescence in CL2070 control animals was arbitrarily normalized as one. The colours highlight individual replicates. One-sample paired t-test was performed. **(D-E)** Animals raised at 25°C exhibit higher *hsp-70* induction upon heat shock than animals grown at 15°C. AM446 animals carrying *rmIs223[hsp-70(C12C8.1)p::GFP]* raised at both temperature were exposed to 30 minutes heat shock at 34°C. Fluorescence was measured three hours post heat shock on a Nikon A1R microscope at 20x magnification. Statistics were performed using a two-way Anova. **(F)** Neuronal stress predicts thermotolerance. PS7171 young adult animals carrying *hsp-16.41p::cGAL/UASp::GFP* were sorted into animals with either 0 to 1 GFP-expressing neurons or >3 GFP-expressing neurons. Animals exhibiting higher levels of neuronal stress exhibit higher thermotolerance. Statistics were performed using a paired t-test. **(G)** Animals expressing GFP under the control of *tax-2* and *tax-4* promoters are not more thermotolerant than wild type (N2) animals not expressing GFP in any neuronal cell (One-way Anova), indicating that the mere expression of GFP in neurons does not provide robustness to stress. Animals were carrying either *muIs164[tax-4p::GFP]* or *sEx13780[tax-2p::GFP]*.

### Variable neuronal stress predicts survival to heat stress

The ectopic over-expression of *hsf-1* in *C.elegans* neurons has been shown to potentiate the heat stress response in peripheral tissues and to extend lifespan by two separable mechanisms (Douglas et al. 2015). To investigate the phenotypic consequences of neuronal stress, we analyzed two independent *C. elegans* lines in which the neural-specific *rab-3* promoter drives *hsf-1* expression in anterior neurons (hereafter referred as *hsf-1^neuro^*, (Douglas et al. 2015)). These two lines express step-wise increments in the levels of *hsf-1*. Indeed, our previous qPCR analysis shows that the *hsf-1^neuro^* line#2 expresses higher levels of *hsf-1* than do either wild type or *hsf-1^neuro^* line#1 (Chauve et al. 2021). Consistent with a dose-sensitive response, the *hsf-1^neuro^* line#2 is the most thermotolerant strain of the two **(Figure 5A**), and its exposure to high temperatures results in a strong induction of *hsp-16.2* in peripheral tissues, as visualized in vivo with the *hsp-16p*::GFP transgene (**Figure 5B-C**).

These experiments reveal that the genetic induction of a heat shock response in neurons is sufficient to protect peripheral tissues from acute heat stress. However, these experiments were done in the context of the genetic over-expression of *hsf-1* above physiological levels. It remains to be shown whether neuronal stress can predict organismal survival under physiological conditions. As shown in **Figure 5I-L**, worms raised at the warmer temperatures (25°C) exhibit higher levels of neuronal stress than worms raised at cooler temperatures. We hypothesized that growth temperature can predict dynamic stress responses. To test this prediction, we have used worms carrying the *hsp-70*(C12C8.1)p::GFP reporter. With this reporter strain, GFP expression is not visible under basal conditions (not shown). We raised those worms either at 25°C or at 15°C, and then exposed adult worms to a short heat shock (30 min at 34°C). As shown **in Figure 5 F-G**, worms raised at 25°C mount a higher heat stress response than worms raised at 15°C. This result suggests that the amount of neuronal stress under basal conditions predicts the extent of the induction of the heat shock response.

To further demonstrate that neuronal stress potentiates the heat stress response, we have used natural variability in neural-induced heat shock responses to determine if the number of neurons with an induced stress response can predict the survival of individual worms to heat stress. To do this, we sorted worms carrying the *hsp-16.41p*::cGAL/UASp::GFP reporter at 20°C, based on the number of GFP activated neurons, and analyzed their survival to a prolonged heat stress. Animals with an already induced stress response in neurons - showing at least 3 GFP activated neurons - exhibit higher levels of GFP (p=0.02) and *hsp-16.2* endogenous mRNA levels (p=0.05) at basal levels (**Figure S6E**). Our results indicate that animals carrying at least 3 GFP expressing neurons survive a lethal stress significantly better than do animals with either 1 or no neurons (**Figure 5F**). The increased robustness of these animals to stress does not depend on a potentially beneficial effect of GFP expression in neurons. Indeed, the over-expression of GFP in *tax-2/4* sensory neurons does not affect the thermal survival of the transgenic animals with respect to GFP deficient strains (**Figure 5G**). Collectively, these results indicate that, under physiological conditions the variability in the induction of neural stress responses has consequences for fitness at the individual worm level, potentially pointing at mechanisms of adaptation under variable environmental conditions.

## DISCUSSSION

In order to monitor inter-individual variability in gene expression for stress response genes at basal level, we developed and validated a new method. qPCR-Bayes applied to “Worm-to-Ct” nanofluidics qRT-PCR is a robust method, validated in vivo, to monitor inter-individual variability in gene expression analysis while accounting for technical variability, even for low-expressed transcripts. The method capitalizes on several elements. First, the Wom-to-Ct cDNA extraction method, which we demonstrate to be more efficient than classical extraction methods (**Figure S1A-B**). Second, the qPCR throughput is increased by nanofluidics technology and the pre-amplification step reduces technical variability. We determined here using *in silico* simulation approaches to determine hyper-parameters for the Bayesian model, and evaluate a reasonable sample size (estimated to 45 single worms) to reliably monitor inter-individual variability (**Figure S2**). Such sample size conveniently allows monitoring of mean and inter-individual variability in two biological conditions in parallel on a nanofluidics chip destined for 96 samples. Lastly, we present a state of the art statistical Bayesian framework called “qPCR_Bayes” destined to uncouple inter-individual variability from technical variability in single worm expression data obtained with the “Worm-to-Ct” coupled with nanofluidics qRT-PCR. Technical variability is monitored by adding synthetic RNA spike-ins during “Worm-to-Ct” lysis part of the method. qPCR_Bayes is an integrated strategy that can perform built-in normalisation using information from all genes, whereas existing qRT-PCR analyses proceed through sequential pre-processing steps to filter and normalize data. This prevents propagation of statistical uncertainty and may affect downstream analyses. We show that qPCR_Bayes, by borrowing information from technical RNA-spike-in performs better than classical normalization using housekeeping genes.

We have validated and calibrated inter-individual variability measured by qPCR_Bayes *in vivo* using transcriptional reporters, thus defining variability reference “gold standards”. It is notoriously difficult to measure inter-individual variability from low-expressed genes due to the known inverse relationship between mean and variability (Eling et al. 2018). Indeed, such intrinsic variability arising from stochastic fluctuations in gene expression dominates for low expressed genes (Stewart-Ornstein, Weissman, and El-Samad 2012). On the contrary, extrinsic variability, caused by variation in the activity of upstream regulators, corresponds to biologically relevant variability. We show that it is possible to capture extrinsic inter-individual variability from low expressed genes. Pairwise correlation on inter-individual expression data for hundreds of stress response genes in the absence of exogenous stress, identified several stress regulons. This is the first study to be able to accurately monitor gene expression from inducible chaperones in the absence of overt stress. We use the inducible *hsps* as “case in point” of regulon from low-expressed genes. Using both smRNA FISH and a highly sensitive bipartite expression reporter for *hsp-16.41*, we validated the measured variability of *hsps in vivo* and we discovered that expression of these *hsps*, under unperturbed conditions, stems mostly from neurons. Further studies will likely elucidate whether neurons have a lower threshold of activation of the heat stress response compared to other tissues, and the mechanisms underlying this.

This study shows that neuronal inter-individual variability in *hsps* expression under basal conditions has phenotypic consequences in isogenic *C. elegans* populations and predicts survival to stress. Work in yeast has shown that variability in gene expression can be adaptive in fluctuating environments with a fitness trade-off between different states of expression (Acar, Mettetal, and Van Oudenaarden 2008). If there is a trade-off between basal *hsp* expression in neurons and another phenotypic trait (such as number of progeny for instance), one could imagine that inter-individual variability in *hsp* expression in neurons might be adaptive at the population level. This is already the case for *hsp-16* expression following heat shock (Casanueva, Burga, and Lehner 2012) and this will require more investigation under basal conditions. Thus, our method paves the way to the study of inter-individual variability in gene expression, as a potentially adaptive biological parameter under fluctuating environments.

The ability to measure inter-individual variability from low expressed genes could be useful for the identification of cancer related genes. There is mounting evidence that changes in variability and overall distribution, missed by differential expression analyses, are particularly important during cancer (Roberts, Catchpoole, and Kennedy 2022). While Bayesian statistical models have been applied to single-cell RNA-sequencing analysis to measure differential mean and variability (Vallejos, Marioni, and Richardson 2015), the high-sensitivity of the Worm-to-Ct method coupled with nanofluidics technology allows better detection of low expressed genes that are undetectable when using single-worm RNA-sequencing (Perez et al. 2017). qPCR_Bayes coupled with high-throughput qRT-PCR could help detect differential variability for low expressed genes in cancer.

## ACKNOWLEDGEMENTS

We are grateful to Dr Christian Lanctôt from BIOCEV in Prague for guiding us and sharing his expertise of single molecule RNA FISH technique in *C. elegans*. We would like to acknowledge Anne Segonds-Pichon from the Bioinformatics facility at the Babraham institute as well as Simon Walker and Hanneke Okkenhaug from the Babraham Imaging facility for their technical help and advice. We thank the CGC (Caenorhabditis Elegans Center) for providing some strains. The CGC is funded by NIH Office of Research Infrastructure Programs (P40 OD010440).

## LIST OF SUPPORTING INFORMATION

### List of supplementary figures

**Figure S1:** Accuracy, sensitivity and specificity of the Worm-to-Ct method.

**Figure S2:** Synthetic data experiments to test the stability of qPCR_Bayes under different hyper-parameter configurations and across different sample sizes.

**Figure S3:** Worm to Ct method coupled with qPCR_Bayes provides more sensitive and reliable single-worm qPCR data than standard methods.

**Figure S4:** Transcriptional reporters used as gold standards faithfully represent endogenous levels of gene expression.

**Figure S5:** Whole body smRNA FISH images and spatial transcriptomics show that inducible *heat shock protein* expression stems primarily from head cells in the absence of heat-shock.

**Figure S6:** Neuronal *hsp* expression at basal level is recapitulated by the bipartite reporter *hsp-16.41p::cGAL/UASp::GFP* and neuronal *hsp* expression is sensitive to temperature.

### List of supplementary tables

**Table S1:** List of target genes monitored using qPCR_Bayes

**Table S2:** List of validated primers used for quantitative RT-PCR in this paper

**Table S3**: Determination of the optimal number of housekeeping genes using geNorm

**Table S4:** Raw Ct values obtained by quantitative Rt-PCR

**Table S5**: Denoised Ct values obtained as output of qPCR Bayes

**Table S6:** Coefficient of variation (CV) of transcriptional reporters

**Table S7**: Number of GFP positive neurons in PS7171 and PS7167

## GENERAL METHODS

### *C. elegans* maintenance

Nematodes were grown on NGM plates seeded with *Escherichia coli* OP50 strain at 20**°**C unless otherwise stated, according to standard methods (Brenner 1974).

### List of *C. elegans* strains used

N2 WT; AGD1054 *hsf-1^neuro^ line #1 uthIs366[rab-3p::hsf-1 full length (FL); myo-2p::tomato]*, AGD1289 *hsf-1^neuro^ line #2 uthIs368[rab-3p::hsf-1 1 full length FL; myo-2p::tomato]*, AM134 *rmIs126[unc-54p::YFP::unc-54 3’UTR],* AM446 *rmIs223[hsp-70(C12C8.1)p::GFP;rol-6(su1006)],* BC13780 *dpy-5(e907)I; sEx13780[rCes F36F2.5(tax-2)::GFP + pCeh361],* BCN1049 *crgIs1004[daf-21p::GFP::unc-54 3’UTR; unc-119(+)],* BCN1050 *crgIs1002[daf-21p::mcherry::unc54 3UTR; unc-119(+)],* BCN1082 *crgSi1004[daf-21p::eGFP)::daf-21 3’UTR]II,* CF1395 *ceh-20(mu290)III*; muIs164[tax-4::GFP], CF2253 *gon-2(q388) ts I,* CL2070 *dvIs70[hsp-16.2p::gfp; rol-6(su1006)]*, GR1395 *mgIs49[mlt-10p::GFP-pest; ttx-1p::GFP] IV,* IG274 *frIs7[nlp-29p::GFP; col-12p::DsRed]*, OH10689 *otIs355 [rab-3p(prom1)::2xNLS::TagRFP] IV*, PS7167 *syIs396 [unc-47p::NLS::NLS::GAL4SK::VP64::let-858 3’UTR + unc-122p::RFP + 1kb DNA ladder (NEB)]; syIs337 [15xUAS::?pes-10::GFP::let-858 3’UTR + ttx-3p::RFP + 1kb DNA ladder(NEB)],* PS7171 *syIs337 [15xUAS::?pes-10::GFP::let-858 3’UTR + ttx-3p::RFP + 1kb DNA ladder(NEB)] III*; *syIs400 [hsp16.41p::NLS::GAL4SK::VP64::let-858 3’UTR + unc-122p::RFP + 1kb DNA ladder(NEB)]*, SS104 *glp-4(bn2) ts*.

We noticed that the strain AGD1289 was prone to silencing and did not exhibit the same appearance and phenotype when grown for more than a month, so we always used freshly thawed AGD1289 worms (less than a month).

### List of *C. elegans* strains generated

MOC1 *mgIs49[mlt-10p::GFP-pest; ttx-1p::GFP] IV x7,* MOC2 *mgIs49[mlt-10p::GFP-pest; ttx-1p::GFP] IV* x7; *glp-4(bn2),* MOC80 *mgIs49[mlt-10p::GFP-pest; ttx-1p::GFP] IV; gon-2(q388) ts I*, MOC86 *crgIs1004[daf-21p::GFP::unc-54 3’UTR; unc-119(+)] x5*, MOC92 *bicIs10[hsp-1p::RFP-AI(artificial introns)::unc-54 3’UTR],* MOC108 *bicSi3[pdaf-21::daf-21::3’UTR unc-54; unc-119(+)] I; unc-119(ed3),* MOC119 *bicls12[ttr-45p::tagRFP-AI(artificial introns)::ttr45UTR],* MOC141 *hsf-1^neuro^line #1 uthIs366 [rab-3p::hsf-1 full length (FL); myo-2p::tomato] x3*, MOC153 *hsf-1^neuro^ line #2 uthIs368[rab-3p::hsf-1 1 full length FL; myo-2p::tomato]; dvIs70[hsp-16.2p::gfp; rol-6(su1006)]*, MOC295 *syIs337 [15xUAS::?pes-10::GFP::let-858 3’UTR + ttx-3p::RFP + 1kb DNA ladder(NEB)] III; syIs400 [hsp16.41p::NLS::GAL4SK::VP64::let-858 3’UTR + unc-122p::RFP + 1kb DNA ladder(NEB)] V; otIs355 [rab-3p(prom1)::2xNLS::TagRFP] IV*.

The genotype of MOC2 worms was determined by checking complete penetrance of the Germline Stem Cell proliferation defect and sterility phenotype after growth at 25**°**C for one generation. The gonadless phenotype of the *gon-2(q388)* ts mutants was not completely penetrant after one generation at 25**°**C. Therefore, *gon-2(q388)* ts animals were grown for two generations at 25**°**C (>90% gonadless phenotype) in our experiments, and the genotype of MOC80 worms was checked by PCR genotyping.

### Worm synchronization for the Worm-to-Ct method

In order to harvest perfectly synchronized individual worms, we used a transcriptional reporter of the molting gene *mlt-10* fused to GFP containing a PEST sequence for rapid degradation (Frand, Russel, and Ruvkun n.d.). The *mlt-10p::GFP-pest* transgene is expressed in the epidermis and fluoresces at the end of every larval stage. At the L4/young adult transition, *mlt-10p::GFP-pest* levels are high and decrease progressively to disappear as worms are become gravid. Individual worms collected for qRT-PCR were first synchronized by egg-lay. Typically, 20 to 30 MOC1 gravid adults carrying *[mlt-10p::GFP-pest]* were transferred onto a NGM plate and removed after 1h. Young adult Day1 animals were harvested about 3h after passing the L4/young adult moult, at 68h post egg-laying synchronization at 20°C, when the levels of fluorescence of *mlt-10p::GFP-pest* were just starting to decrease after the L4/young adult molt. For experiments with *glp-4* mutants, worms were maintained at 15°C and switched to 25°C for the experiment, in order to obtain the germline stem cell proliferation phenotype. For experiments with *gon-2(q388)* temperature sensitive mutants, worms were maintained at 15°C and switched for two generations at 25°C in order to obtain full penetrance of the gonadless phenotype. For worm synchronization in the absence of the *mlt-10p::GFP-pest* transgene, animals were age-synchronized either by egg-lay in a 2h period or by treatment with alkaline hypochlorite solution for experiments that required large amounts of synchronized animals (smRNA FISH, PS7171 sorting). For bulk qRT-PCR experiments and microscopy quantification of fluorescent reporters on L4.8 or L4.9 synchronized worms (Mok, Sternberg, and Inoue 2015) grown in parallel at different temperatures, worms were harvested 97h after egg-lay at 16**°**C, 42h after egg-lay at 25**°**C, and L4.8 or L4.9 worms were picked from a mixed population grown at 20**°**C.

### Single-worm sample preparation by the Worm-to-Ct method

We developped and optimised a protocol to perform quantitative qRT-PCR on single worms, which we call “Worm-to-Ct protocol”. This protocol is based on Power SYBR® Green Cells-to-CT™ kit and downstream high-throughput quantitative RT-PCR using nanofluidic technology using a Biomark®, We have previously published a detailed protocol to perform quantitative RT-PCR on single-worms (Chauve et al. 2020). The Biomark® HD system using nano-fluidic chips allows high-throughput qRT-PCR. The 96.96 Gene Expression array produces 9216 qRT-PCR results in a single run (96 different couples of primers run on 96 single worms). We also used smaller chips, the Flexsix IFC, whose set up was flexible: either 12 target genes x 72 samples, or 24 target genes x 36 samples).We recapitulate here the main steps and add details specific to the use of Worm-to-Ct protocol coupled with the statistical model qPCR_Bayes, such as adding synthetic RNAs and data filtering. In brief, single worms are directly added to the lysis buffer, skipping direct isolation of easily degradable RNA. 12 technical spike-ins (synthetic RNA transcript of known sequence and quantity from ERCC) spanning a broad range of concentrations are added to the lysis buffer. RNA capture is boosted by 10 freeze-thaw cycles and a 20-minute vortex step to break the worm’s cuticle. The obtained lysed worm containing RNA is reverse transcribed and subject to a pre-amplification step. Primers are designed across introns to ensure amplification of only spliced transcripts (**Figure S6A**). Primers for qPCR are validated by using a standard curve and only those showing high specificity and PCR efficiency between 90 and 115% are further used for downstream analysis (Table S1). qPCR is run on a 96.96 nanofluidics Chip from Fluidigm. In each Chip, 96 single worms and 96 primer sets which result in up to 9216 reactions per experiment and is run using the Biomark®. In one experiment, 81primers are used to amplify targets, 3 primers to amplify house-keeping genes used for normalisation of data and the remaining 12 primers are used to amplify the exogenously added ERCC spikes (**Figure S6A**). Based on simulation experiments on a set dataset comprising 90 single worms (**Figure S9 B-E**), we have estimated that 47 single worms are enough to estimate inter-individual variability reliably and therefore two conditions can be directly compared in 1 single Chip. The raw data obtained from Biomark is filtered in two ways. First, a manual step is used to eliminate all failed reactions and second, only Ct values that surpass the estimated limit of detection (LOD>22) are kept numeric (**Figure S6B**), values that are lower than the Ct (LOD< = 22) are substituted for (Not a number: NaN values). This filtered raw data is then processed using the qPCR Bayes pipeline.

### Quantitative RT-PCR on bulk worm samples using the Worm-to-Ct method

To monitor steady-state mRNA levels on bulk samples of worms, we used the Worm-to-Ct, based on the Power SYBR® Green Cells-to-C_T_™ kit and downstream quantitative RT-PCR using nanofluidic chip on a Biomark®, with a few adaptations. We hand-picked a pool of about 15-30 worms in 10 µL Lysis buffer. Reverse transcription was performed using the 2X RT buffer from the Power SYBR® Green Cells-to-C_T_™ kit according to manufacturer instructions and cDNA was diluted either 1:4 or 1:5. Each qRT-PCR reaction contained 1.5 µL of primer mix forward and reverse at 1.6 µM each, 3.5 µL. The full protocol is detailed in (Chauve et al. 2020).

### Extraction of RNA from single worms with Trizol

Individual worms were collected in RNase free water to which 500 µLof Trizol Reagent (Invitrogen, 11596018) and 1µl Glycoblue Coprecipitant (Invitrogen, AM9515) were added and tube flash frozen in liquid nitrogen. RNA was extracted as per (He 2011). RNA concentration and quality were determined by running on an Agilent Bioanalyzer using an RNA 6000 Pico kit (Agilent, 5067-1513). Samples with a RIN of 9 or higher were used for further analysis.

### Quantification of primer efficiency and specificity

Primers sets were designed to quantify *C. elegans* post-spliced transcripts (listed in Table S1). Primer sets were designed to span exon-exon junctions using NCBI Primer Blast software and subsequently blasted against the *C. elegans* genome to test for off-target complementarity. The first method used was standard Trizol extraction and reverse transcription using SuperScript IV. The second, method used the Worm-to-Ct protocol (see above) on bulk samples to generate the cDNA. The cDNA samples were diluted in a series of 4 to 6 times. Each qRT-PCR reaction contained 1.5 µL of primer mix forward and reverse at 1.6 µM each, 3.5 µL of nuclease free water, 6 µL of 2X Platinum® SYBR® Green qPCR Supermix-UDG (Thermo Fisher Scientific PN 11744-500) and 1 µL of worm DNA lysate diluted or not. The qRT-PCR reactions were run on an iCycler system (Bio-Rad).

### 2.7-Choice of ERCC RNA spike-in RNAs to monitor technical noise

The ERCC RNA spike-in mix (ThermoFisher PN4456740) comprises a mix of 92 external polyA transcripts developed by the External RNA Control Consortium (ERCC). The synthetic RNA spike-in are 500bp up to 2000 bp and cover 8000X range concentration. A set of 12 ERCC RNAs spike-in (ERCC#9, ERCC#25, ERCC#31, ERCC#34, ERCC#51, ERCC#60, ERCC#67, ERCC#126, ERCC#148, ERCC#150, ERCC#158, ERCC#171), of various concentrations, was used to monitor technical noise on every 96.96 nano-fluidic chip. The ERCCC kits contains two mixes (mix 1 and mix 2) with some RNA spike-in spike-ins at different concentration. Our chosen set of 12 ERCC RNAs is at the same concentration in mix 1 or mix 2, so both mixes could be indifferently used. When using the small nano-fluidic FlexSix IFCs in the configuration, 12 target genes x 72 single worms, a set of 3 ERCC RNA spike-in (ERCC#25, ERCC#51, ERCC#60) was used.

### Filtering and normalisation of raw data

First the data was pre-processed by eliminating columns (containing raw data per assay) or rows (containing raw data per individual worms) where no values were detected. Second, undetected reactions (Raw output=999) were redefined as “not a number” (NaN) values. Values that fall below the estimated LOD (raw Ct values equal or higher than 22) were also replaced by NaN values. Finally, columns of data that had more than 20% NaNs were filtered. All NaN values were imputed to the LOD value of 22. To obtain delta Ct values, first, the R code calculates the geometric mean of normalising genes (*ire-1, cdc-42 and pmp-3*) per row. Second the delta-Ct values are calculated by subtracting the geometric mean to each raw Ct value for the same sample. In this way, every assay per worm is normalised.

### Worm DNA lysates preparation and PCR genotyping

A few worms were picked into 10 µL of Worm Lysis Buffer (50mM KCl, 10mM Tris (pH 8.3), and 2.5mM MgCl2, 0.45% NP40, 0.45% Tween-20, 0.01% Gelatin). Tubes were freezed-thawed 10 times before adding 1 µL 01mg/mL of proteinase K. Worms were lysed and genomic DNA was released by heating tubes to 65**°**C for 60-90 minutes. Proteinase K was inactivated by heating to 95**°**C for 15 min. Commonly, 1 μL DNA lysate was added to PCR reaction. All PCR genotyping reactions for genotyping were performed with Taq DNA polymerase (New England Biolabs #M0267L) with Thermopol buffer according to manufacturer instructions. The *gon-2(q388)* mutation is a G->A change at nucleotide 2878. A 752 bp PCR product was amplified using the primers below and the purified PCR product was sequenced. PCR conditions were: 95**°**C 30s, 35 cycles: 95**°**C 20s, 61**°**C 30s, 68**°**C 60s, and 68**°**C 5 min. The sequence of *glp-4(bn2)* is not available. Therefore *glp-4(bn2)* were identified based on sterility phenotype at 25°C. Sequence of the primers used for PCR genotyping:

gon-2(q388) For 5’-GCTACCAACGCTTTTGCCAT

gon-2(q388) Rev 5’-AGACAAGTATAAACAACGGATGAAA

### Cloning and Transgenics

**Table 2:**
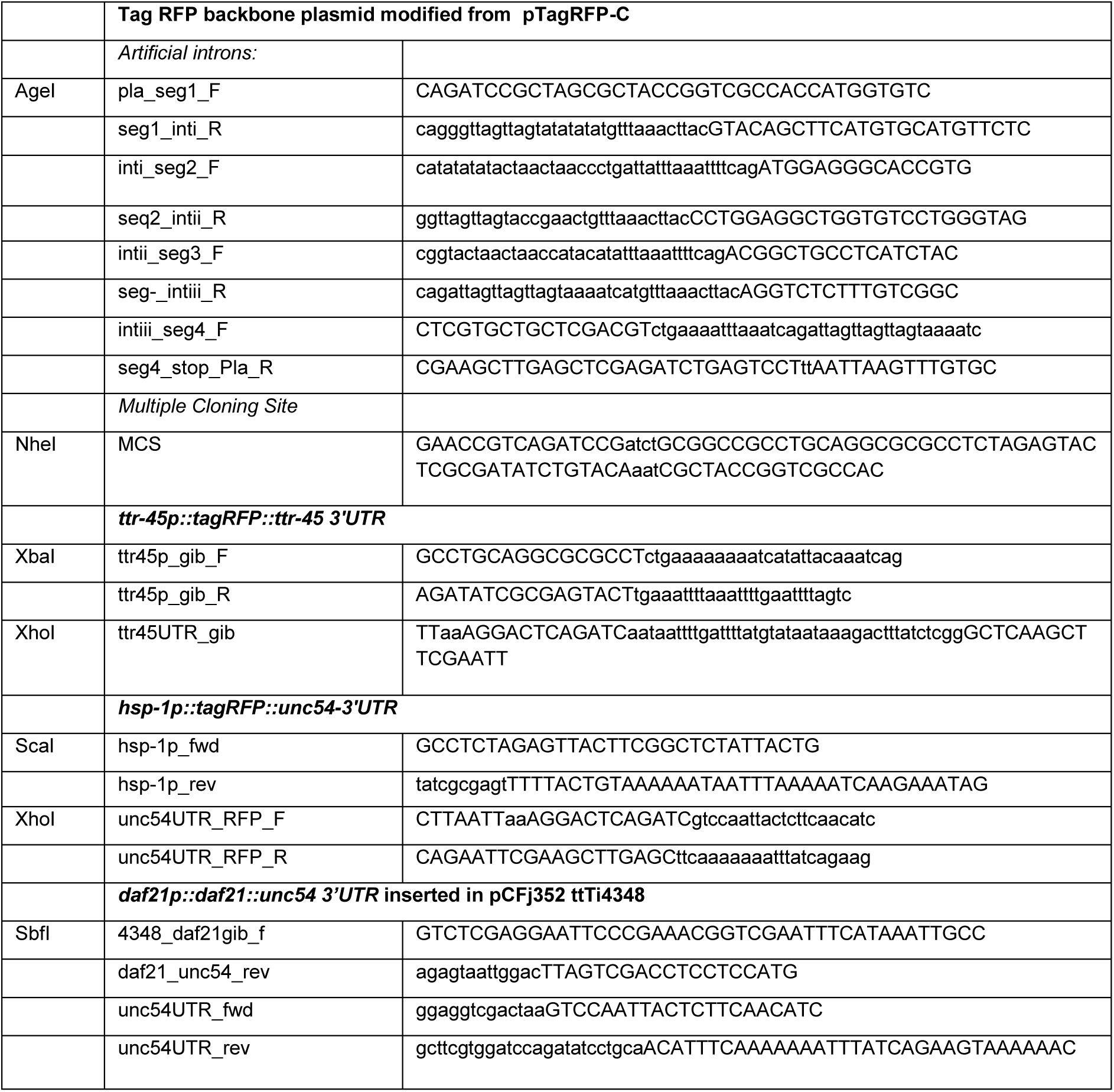
Primers used for cloning.

All cloning was performed using Gibson Assembly Master mix (New England Biolabs, E2611) as per manufacturer’s protocol and the listed primers. The pTagRFP-C (Evrogen) plasmid was modified to add *C.elegans* artificial introns and an N-terminal multiple cloning site; this plasmid was used for subsequent TagRFP constructs. Injections were performed into N2 worms as per standard protocols using pCFJ421 (Addgene) as co-injection maker. The *daf21p::daf21::unc54 3’UTR* construct was cloned into pCFJ352 (Addgene) and injected into EG6701 as per http://www.wormbuilder.orgprotocol.

### Strain integrations

Extra-chromosomal strains MOC92 and MOC119 were integrated by UV irradiation according to the fast integration protocol developed by (Mariol et al. 2013). A Stratagene Stratalinker UV cross linker was used with a dose of energy ranging from 30 to 35 mJ/cm^2^.

### Heat-shock

NGM plates were double sealed with parafilm and heat shocked in a water bath. Unless otherwise stated, heat shock was 30 minutes at 34**°**C. Worms were typically harvested immediately after heat shock.

### Thermotolerance assay

The thermotolerance assay was performed as described previously (Labbadia and Morimoto 2015), except that heat-shock was 4h at 36**°**C. About 60-100 synchronized animals were picked onto a 6 cm NGM plate with 2 plates per condition. For PS7171, animals were sorted at the young adult stage. For N2, BC1378 and CF1395, thermotolerance was performed at day 2 of adulthood, defined as 24h post L4.8 stage. Animals were then allowed to recover overnight at 20**°**C. Animals were scored the next day at about 20h after the end of heat-shock. Animals were transferred onto a new plate, and were counted as alive when they were either moving on the plate, or at least able to move their nose when poked with a pick.

### 3.8-Single molecule RNA FISH (smRNA FISH)

The probes were designed using the Stellaris Probe Designer software (Biosearch technologies): https://www.biosearchtech.com/support/tools/design-software/stellaris-probe-designer. We used Qasar 570 for *hsp-70* and *hsp-16* probes and Qasar 670 for *hsp-1* probe. 48 probes were designed for *hsp-1* and *hsp-70 (C12C8.1)* based on the spliced ORF sequence and using default settings using. However, as the *hsp-16.41* gene was too small to allow the design of enough probes, we included 5’UTR and 3’UTR elements and lowered the masking levels to 0, allowing to create 27 probes (minimum=25). Each probe was individually blasted against the *C. elegans* genome. While the *hsp-16* probe likely hybridizes against other members of the *hsp-16* family, it does not appear to match other mRNAs. SmRNA FISH was performed according to the protocol developed by (Bolková and Lanctôt 2016), using custom-based worm baskets to transfer the worms and 3h hybridization at 37**°**C. About 1000 of young adults synchronized by hypochlorite treatment were harvested per condition. As a control, we added heat shock worms in every experiment.

### 3.9-Microscopy

All worms imaged were mounted on a 2% agarose pad. For imaging of live worms, animals were paralyzed in 3mM Levamisole diluted in M9. Fluorescence exposure was identical across all conditions of the same experiment. For imaging GFP positive neurons in PS7171 and PS7167 we used a Nikon fluorescent stereomicroscope SMZ18, as it was easier to capture neurons in 2D from live animals. PS7171 and PS7167 were synchronized at L4.8 stage (Mok, Sternberg, and Inoue 2015) in this experiment. To determine the levels of fluorescence in synchronized worms of our gold standard transcriptional reporters, MOC92, MOC119, and AM134 **(Figure 1**) were imaged on the Nikon SMZ18, while IG274 (**Figure 1**), MOC86, BCN1049 and BCN1050 (**Figure S1**) worms were imaged on the Nikon Ti Eclipse at objective 20X objective. MOC295 (**Figure 3G-H**) and AM446 (**Figure 5D**) animals were also imaged on Nikon Ti Eclipse at objective 20X objective. Worms were prepared for smRNA FISH (**Figure 3A-D**, **Figure 4G-H, Figure S4A-H**) were mounted in prolong Gold and imaged on the Nikon A1R confocal microscope at 63X objective. The Z-stacks taken were then processed by deconvolution and stitched together.

