## Supplementary material for "Inter individual variability in neuronal expression of heat shock protein genes predicts stress survival in *Caenorhabditis elegans*": Suplemental Figures

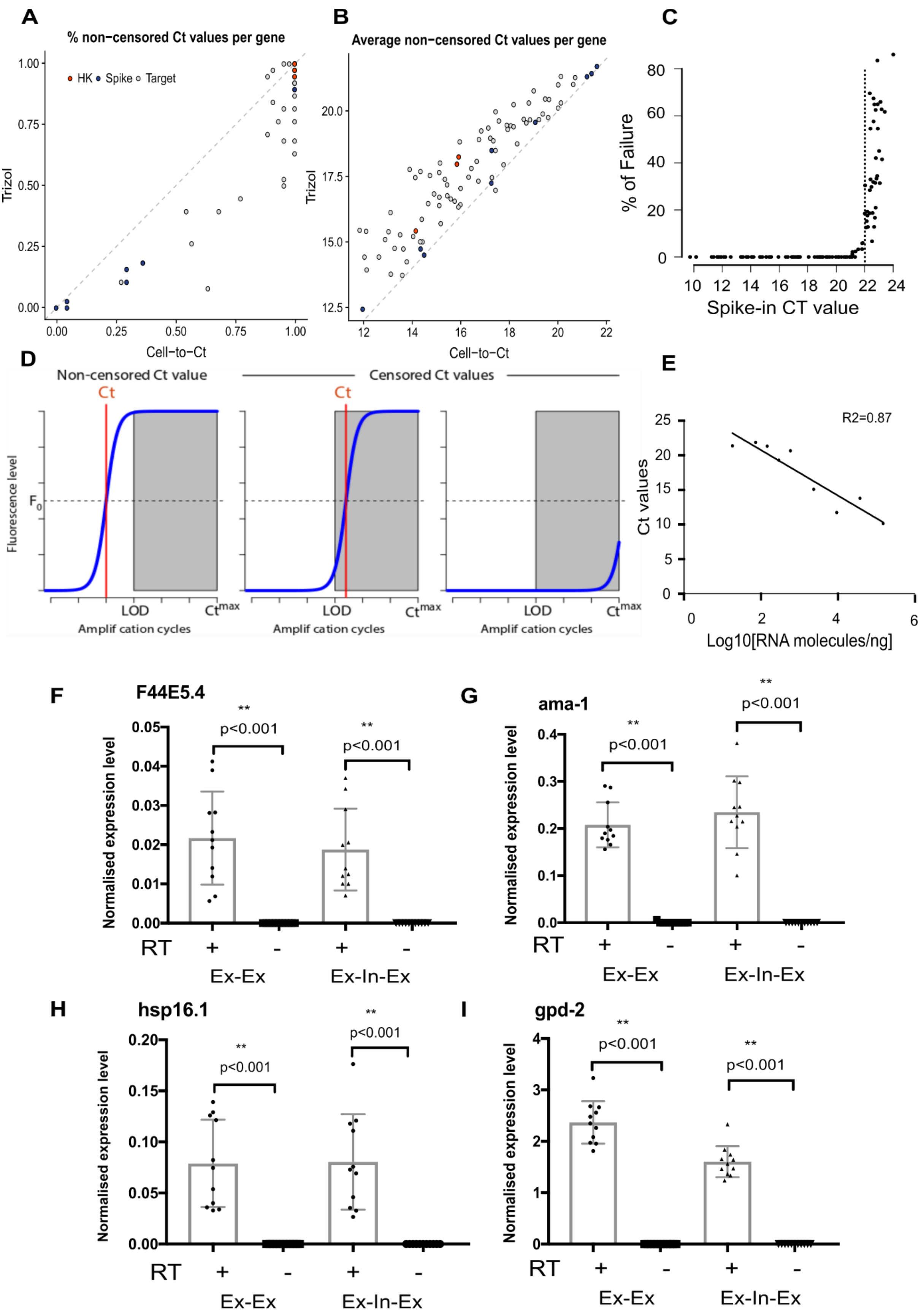
A
% non−censored Ct values per gene
HK
Spike
Target
Trizol
Cell−to−Ct
1.00
0.75
0.50
0.25
0.00
B
Average non−censored Ct values per gene
Trizol
Cell−to−Ct
20.0
17.5
15.0
12.5
12
14
16
18
20
22
C
% of Failure
Spike-in CT value
80
60
40
20
0
10
12
14
16
18
20
22
24
D
Non-censored Ct value
Censored Ct values
Ct
Fluorescence level
F0
LOD
Ctmax
Amplif cation cycles
E
R2=0.87
Ct values
Log10[RNA molecules/ng]
25
20
15
10
5
0
2
4
6
F
F44E5.4
Normalised expression level
0.05
0.04
0.03
0.02
0.01
0.00
**
p<0.001
RT
+
-
Ex-Ex
Ex-In-Ex
G
ama-1
0.4
0.3
0.2
0.1
0.0
H
hsp16.1
0.20
0.15
0.10
0.05
0.00
I
gpd-2
4
3
2
1
0

**Figure S1. Accuracy, sensitivity and specificity of the Worm-to-Ct method. Worm to Ct method coupled with qPCR_Bayes provides more sensitive and reliable single-worm qPCR data than standard methods.** (**A-B**) Comparison between the Trizol and Worm-to-Ct extraction methods applied to the same group of day 1 of adulthood worms grown in parallel (plate 21, tables S2 and S3). (**A**) Percentage of non-censored observations for each gene. In this calculation, censored observations are defined as those of which the observed Ct value exceeds the limit of detection (set equal to 22, Figure S1A). (**B**) For each target gene, Ct values obtained with Worm-to-Ct are overall lower than Ct obtained with Trizol, indicating higher expression levels. Across all target genes, Ct were on average 1.43 Ct lower with Worm-to-Ct compared to Trizol, indicating that cDNA levels are on average 2-4 fold higher using Worm-to-Ct than Trizol. For each gene, mean of the observed Ct value across all non-censored observations. Only genes with at least two non-censored observations are displayed. (**C**) Estimation of sensitivity expressed as LOD (Limit Of Detection) using ERCC RNA spike-ins. To estimate LOD we added synthetic RNA spike-in to the lysis buffer. Synthetic RNA of different lengths (between 500 and 2000 nucleotides) were used at concentrations covering a broad dynamic range (from 0.4578 to 3750 attomoles/μL). The figure shows the % of failure as a function of the average Ct values for the set of 12 RNA spike-in monitored (data from all plates). The 5% failed PCR cut off corresponds to a Ct value of ~22. Any Ct>22 value obtained in our experiments is thus considered not reliable. (**D**) Censoring in qRT-PCR gene expression assays. In qRT-PCR assays, $Ct$ values are defined as the number of PCR amplification cycles that were required to achieve a given fluorescence level threshold $F_0$ (left panel). Censoring arises when actual $Ct$ values are "undetermined". Two cases can occur: $F_0$ is achieved but the observed $Ct$ value exceeds the limit of detection (LOD) above which values are deemed unreliable (middle panel), or when the fluorescence threshold $F_0$ was not achieved (right panel). (**E**) Estimation of the accuracy of the method by a linear correlation between raw Ct values and known concentration of synthetic ERCC RNA spike-in. The concentration of the pre-amplified synthetic RNAs spike-in in number of molecules per ng of RNA is provided by ERCC and sold by ThermoFisher (PN4456740). The linear regression shows a high correlation between the two values ($R^2$=0.87, p<0.0002, F-test). The lowest amount detected falls below 600 molecules per μl (< 200 copies/ng of total RNA, data from plate 15). We considered Ct value of 22 as a threshold above which the data is not reliable. (**F-I**) Worm-to-Ct method for sample preparation effectively eliminates genomic DNA contamination. The graph shows the expression level per worm and the standard deviation of quantitative RT PCR reactions obtained from Worm-to-Ct cDNA for 4 target genes. Targets were normalised to *cdc-42* and transformed to natural scale by the delta Ct method and include (F) *F44E5.4,* (G) *ama-1,* (H) *hsp16.1,* (I) *gpd-2*, where cDNA has been subject to reverse transcription (+RT) or not (-RT). The first set of primers, designed exclusively in exons, (Ex-Ex) can anneal only to spliced mRNA and not to genomic DNA (gDNA). A second set of primers (Ex-In-Ex) were designed complementary to exons flanking a short intron (intron size <100 bp). Ex-In-Ex primers are therefore able to amplify from gDNA template as well as from cDNA coming from both nuclear unspliced transcripts as well as mature mRNA. The graph reveals that there is no amplification in the -RT samples (p<0.01, t-test), where Ex-In-Ex primers should be able to amplify gDNA, indicating the absence of genomic DNA contamination in single worm lysates using the Worm-to-Ct protocol.

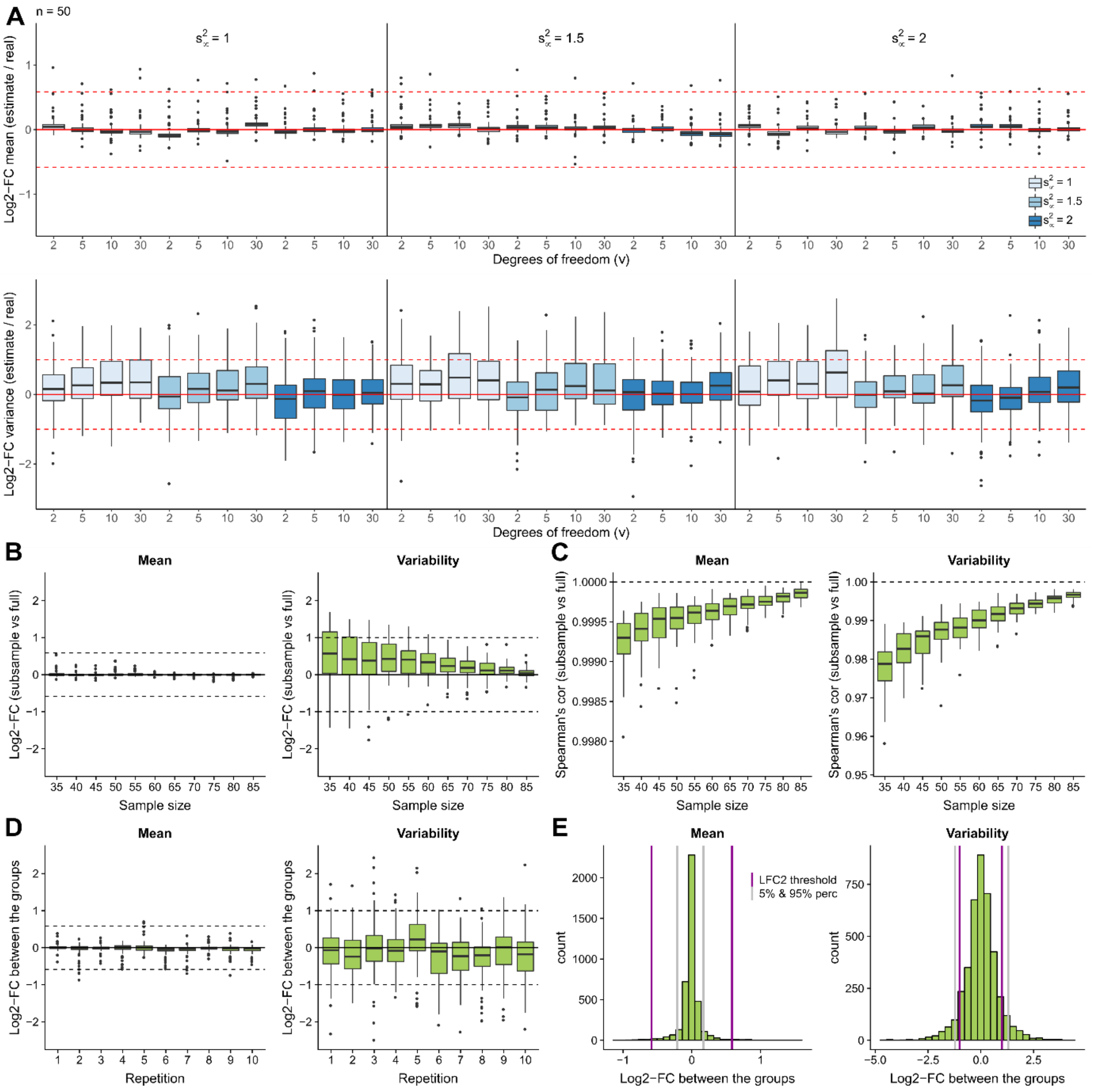


**Figure S2: Synthetic data experiments to test the stability of qPCR_Bayes under different hyper-parameter configurations and across different sample sizes.** (**A**) Synthetic data experiments were performed to assess the performance of qPCR_Bayes under different hyper-parameter configurations. Synthetic data (n=50 samples) was generated using the statistical model implemented in qPCR_Bayes. True parameter values were set to match the empirical estimates obtained for the plate 7 dataset (tables S2 and S3). For each hyper-parameter configuration, the log2 fold change (log2-FC) between parameter estimates and their associated true values was calculated. Boxplots summarize the distribution of these log2-FCs across all genes for mean expression (upper panel) and variability (lower panel) parameters. Based on these results, default hyper-parameter values were set to s2μ=2, s2δ=2 and v=5. (**B-C**) Based on the plate 7 dataset (90 samples from a single experimental condition) a random sub-sampling strategy was used to assess the stability of qPCR_Bayes across different sample sizes. For each sample size, 50 repetitions were run using different random sub-samples of the data. Parameter estimates based on the full dataset were used as a benchmark to evaluate estimation performance for smaller sample sizes. (**B**) For a single repetition, boxplots summarize the distribution (across genes) of the log2-FCs between the estimates based on the sub-sampled and the full data for mean expression (left panel) and variability (right panel) parameters. (**C**) Boxplots summarize the distribution (across all 50 repetitions) of the Spearman’s rank correlation between estimates based on the sub-sampled and the full data for mean expression (left panel) and variability (right panel) parameters. (**D-E**) Based on the plate 7 dataset (90 samples from a single experimental condition) a random data partition experiment was used to generate two groups of samples (n=45 each, matching our experimental design) for which we do not expect to detect any differentially expressed genes. This enables us to assess the range of log2-FCs that can be expected by chance for a limited sample size. (**D**) For 10 repetitions of our data partition experiment, boxplots summarize the distribution of the estimated log2-FCs between the groups for mean expression (left panel) and variability (right panel) parameters. (**E**) Pooling the results obtained across 50 repetitions and across all genes, distribution of the estimated log2-FCs between the groups for mean expression (left panel) and variability (right panel) parameters. Vertical grey lines denote the 5% and 95% percentiles of the log2-FC distributions. Vertical purple lines denote our chosen default log2-FC thresholds.

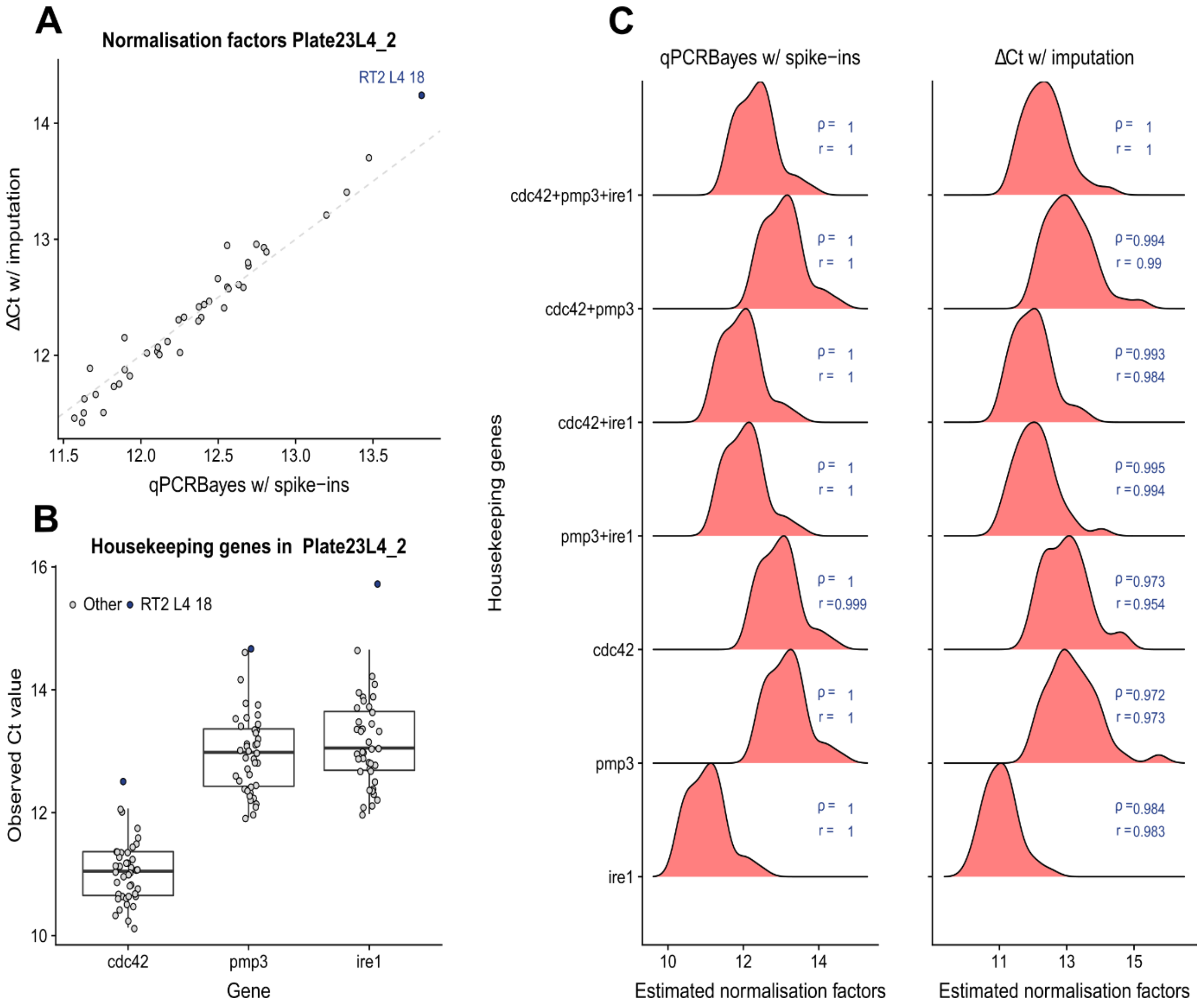


**Figure S3: Two strategies for normalization when using Worm to Ct method coupled with qPCR_Bayes.** (**A**) Comparison between estimated normalization parameters estimated by qPCR Bayes (spike-ins version) and the Δ Ct method. This comparison is based on the data produced in plate 23, second group of samples (tables S2 and S3). (**B**) For the second group of samples in plate 23, distribution of the observed Ct values of the housekeeping (HK) genes across all samples. (**C**) Distribution of the estimated normalization factors when using different sets of HK genes. Pearson's (ρ) and Spearman's (r) correlations with respect to the estimates obtained when using all three genes as housekeeping genes (top row) are provided.

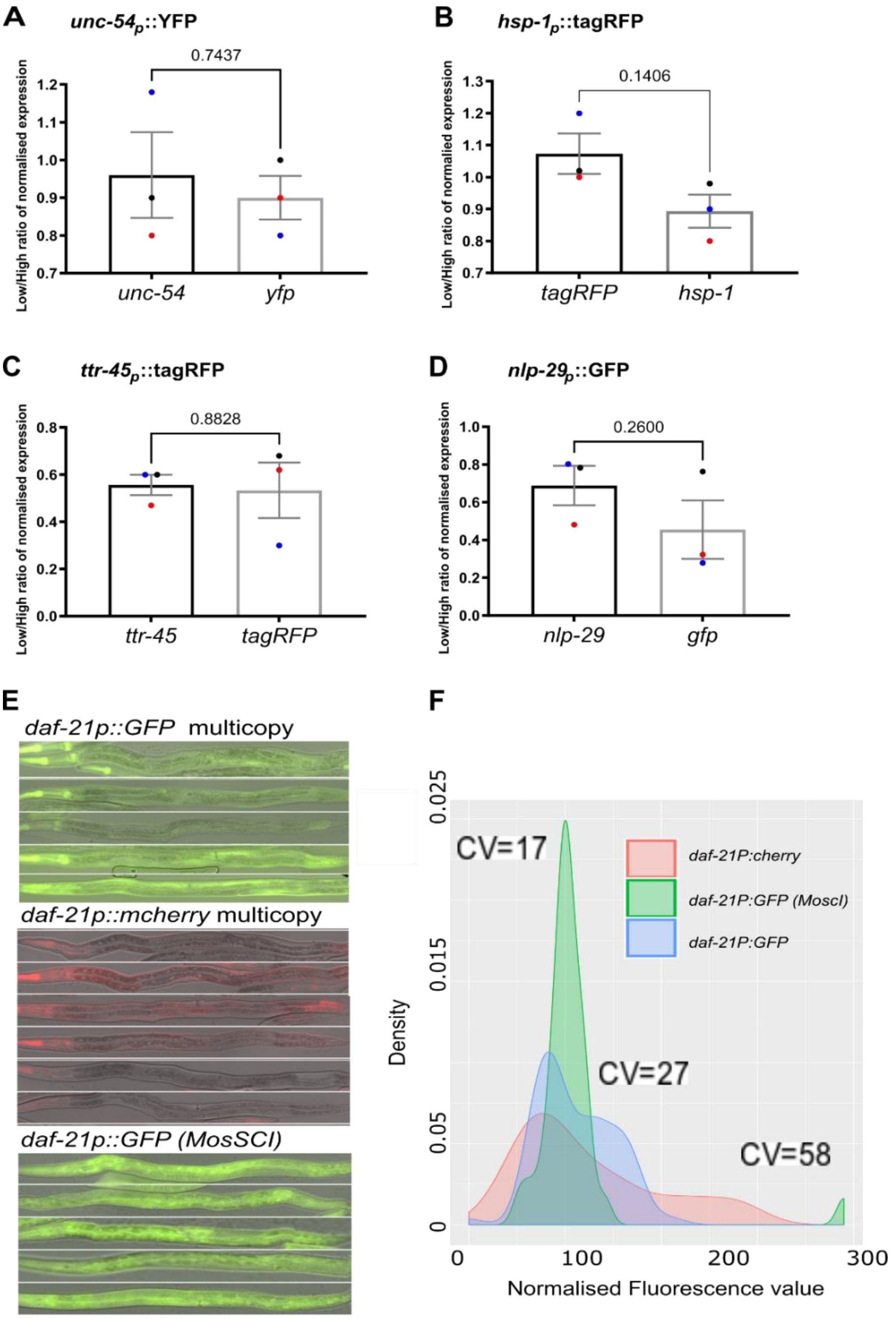


### G. Correspondence between the transgene and the endogenous locus

| | *unc-54p:YFP* | *hsp-1p:tagRFP* | *ttr-45p:tagRFP* | *nlp29p:GFP* |
|---|---|---|---|---|
| Average ratio low/high fluorophore (+/- SEM) | 0.96 +/- 0.11 n=3 | 0.89 +/- 0.05 n=3 | 0.55 +/- 0.04 n=3 | 0.689 +/- 0.1 n=3 |
| Average ratio low/high gene (+/- SEM) | 0.9 +/- 0.05 n=3 | 1.073 +/- 0.06 n=3 | 0.53 +/- 0.1 n=3 | 0.455+/- 0.15 n=3 |
| P value (paired t-test) | 0.74 | 0.14 | 0.88 | 0.26 |
| Significance | No | No | No | No |

**Figure S4. Transcriptional reporters used as gold standards faithfully represent endogenous levels of gene expression.** (**A-D**) Gold standards (GS) are used to measure inter-individual variability, it is therefore key that the variability captured by the GS is close to that of the endogenous transcripts. To test this relationship, we compared the ratio of mRNA obtained from animals sorted as low versus high for all GS transgenes. The data represents the ratio of relative expression level (and standard error) of low versus high groups, for either endogenous or fluorophore mRNA. In vivo transcriptional reporters (all multi-copy integrated arrays) for (**A**) *bicIs10[hsp-1p::RFPAI::unc-54 3'UTR]*, (**B**) *rmls126 [unc-54p::YFP::unc54 3'UTR],* (**C**) *bicls12[ttr-45p::tagRFP::ttr453'UTR]*, (**D**) *frIs7 [nlp-29p::GFP + col-12p::DsRed]*, were sorted based on levels of fluorescence into high and low expressors and processed for standard bulk qRT PCR. The timings correspond to those used in Table S4. All genes were normalised to the average of three housekeeping genes (*cdc-42, pmp-3* and *ire-1*) and relative transcript expression was quantified using standard bulk qPCR normalised using delta Ct method as described in materials and methods. The dots have been colour coded by replicate pairs**.** Statistics: t-test on three biological replicates. Statistics and detailed values for comparisons between ratios are depicted in G. (**E-F**) In our studies, we noticed that a transcriptional reporter of the *hsp-90/daf-21* promoter is highly variably expressed when integrated in multiple copies in the genomes, referred to as *daf-21p::GFP* and daf-*21p::mcherry* in (E,F), but not when integrated as a single copy (referred to as *daf-21p::mosSCI* in (E,F). The single copy transgene that drives stable gene expression contains an endogenous *daf-21* 3'UTR whereas the multi-copy transgenes contain the heterologous *unc-54* 3'UTR. To determine if the variability of the transgenic animals was related to the exogenous 3'UTRs, we constructed a single copy insertion of *daf-21* promoter using *unc-54* as 3'UTR: *bicSi3 [daf-21p::daf-21::unc54::3'UTR]* to parallel multicopy transgenes, and measured biological variability using qPCR_Bayes (Figure 4D). However, the variability of the *bicSi3* single copy transgene is very low (Figure 4D) and similar to endogenous *daf-21* transcript (table S3), indicating that the multi-copy nature of the inserted transgenes and not their heterologous 3'UTR sequences causes their increased variability. (**E**) Representative images of *hsp-90/daf-21* reporters: BCN150: *crgIs1002 [pdaf-21p::mcherry::unc543UTR],* MOC86: *crgls1004 [daf-21p::GFP::unc543'UTR]* and MOC85 *crgSi?[daf-21p::egfp::daf-21 3'UTR].* (**F**) Density distribution of multicopy *hsp-90/daf-21* reporters (*daf-21p: mcherry* and *daf-21p:GFP*) and single copy-integration (*daf-21p:GFP(MoscI).* Additional data in Table S4. (**G**) The table shows the values for paired t-test statistics on the proportion of change between high and low groups for either fluorophore (first row) or endogenous (second row) transcripts. This analysis indicates that reporters are faithfully capturing the variability of the endogenous gene.

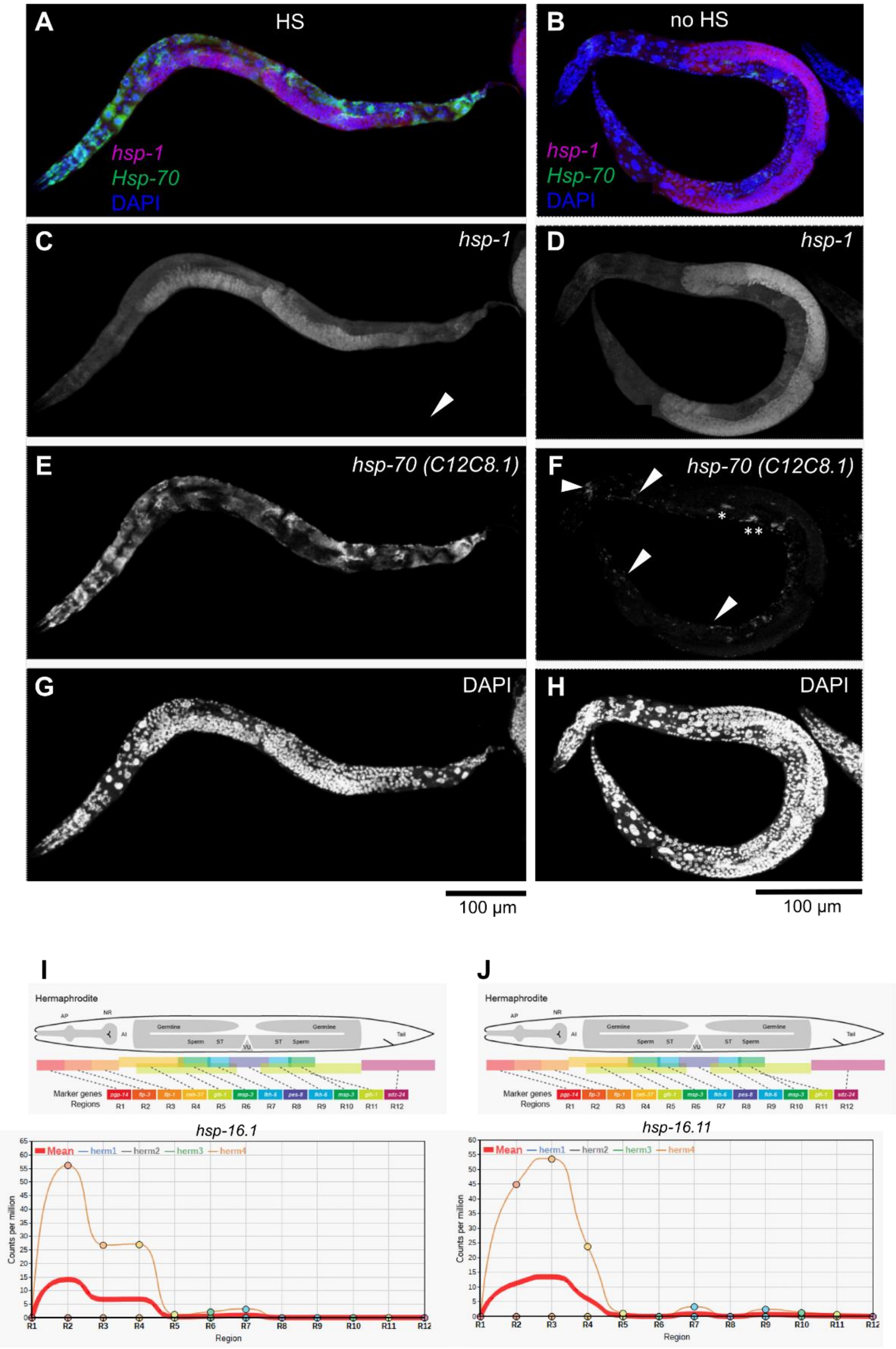
A
HS
hsp-1
Hsp-70
DAPI
B
no HS
hsp-1
Hsp-70
DAPI
C
hsp-1
D
hsp-1
E
hsp-70 (C12C8.1)
F
hsp-70 (C12C8.1)
*
**
G
DAPI
H
DAPI
100 µm
100 µm
I
Hermaphrodite
Marker genes
Regions
R1 R2 R3 R4 R5 R6 R7 R8 R9 R10 R11 R12
hsp-16.1
Mean — herm1 — herm2 — herm3 — herm4
Counts per million
Region
J
Hermaphrodite
Marker genes
Regions
R1 R2 R3 R4 R5 R6 R7 R8 R9 R10 R11 R12
hsp-16.11
Mean — herm1 — herm2 — herm3 — herm4
Counts per million
Region

**Figure S5. Whole body smRNA FISH images and spatial transcriptomics show that inducible *heat shock protein* expression stems primarily from head cells in the absence of heat-shock. (A-H)** Fluorescent micrographs (maximum intensity projection) of single molecule RNA Fluorescent in-situ hybridization (smRNA FISH) using probes against inducible *hsp*-70(C12C8.1) (*ihsp70*), in green (E,F), constitutive molecular chaperone *hsp-1*, in red (C,D) and DAPI (G,H) to stain nuclei in blue. FISH was performed on WT day1 young adult animals after a 30 minutes heat shock at 34°C (**A,C,E,G**) or in the absence of heat shock **(B,D,F,H**). Note that after heat shock *ihsp70* transcripts are present in most somatic tissues at high levels. In the absence of heat shock, *ihsp-70* transcripts were present in head cells (two asterisks) and in some animals in somatic gonadal cells (one asterisk). In contrast, transcripts for constitutive *hsp-1* are present in most cells with and without heat shock, and are strongly enriched in the gonad. Scale bar represents 100 µm. Images were taken with 60x objective using a Nikon A1R confocal microscope. (**I-J**) Graphs depicting the results of spatial transcriptomics analysis (Ebbing et al. 2018) in adult hermaphrodites for *hsp16.11* and *hsp16.1* mRNAs*.* Transcripts of *hsp-16.1* and *hsp-16.11* were both enriched in the head region of the worm. Upper panel graphically shows the regions that were assayed (Images were downloaded from: *http://celegans.tomoseq.genomes.nl).*

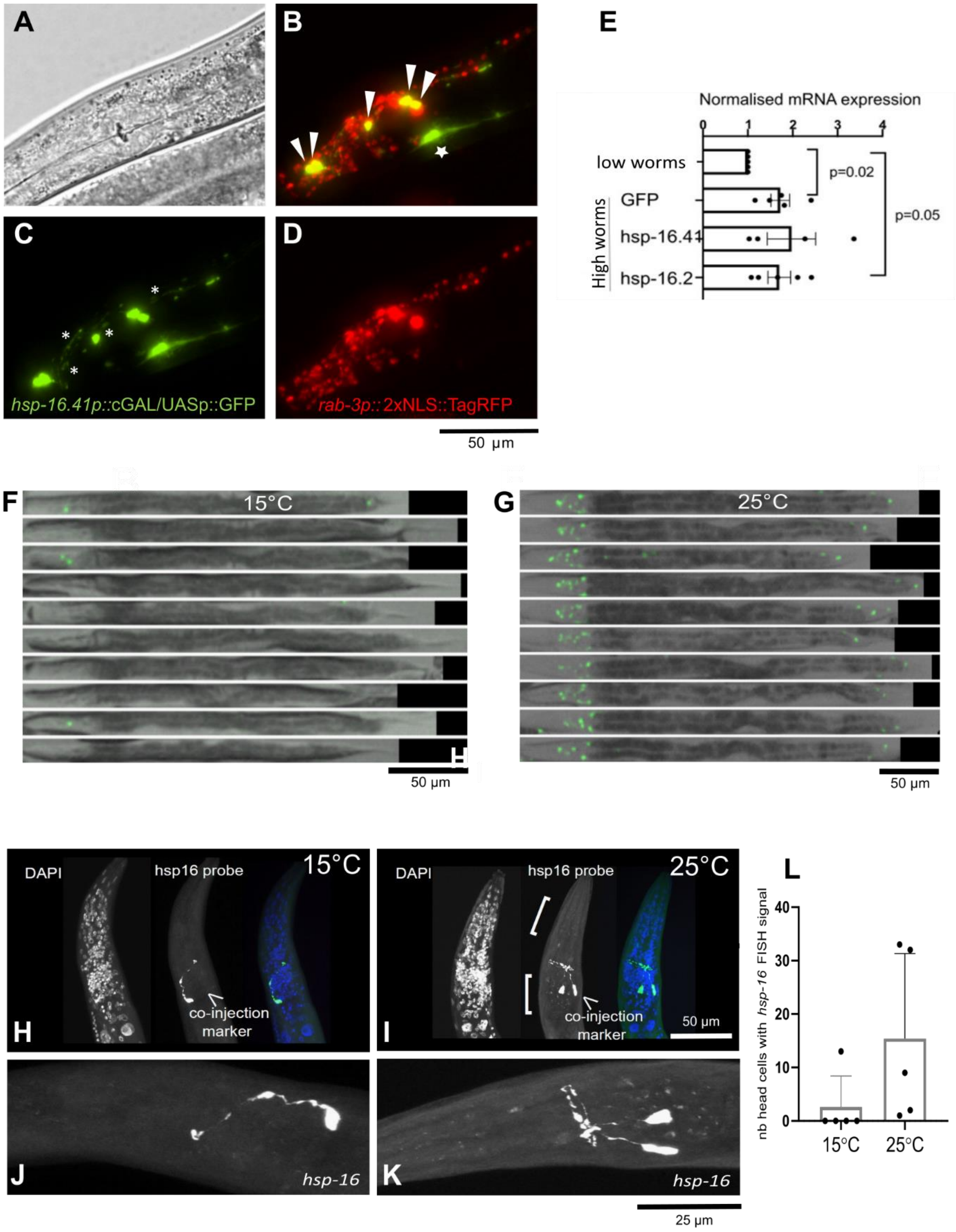


**Figure S6. Neuronal *hsp* expression at basal level is recapitulated by the bipartite reporter *hsp-16.41p::cGAL/UASp::GFP* and neuronal *hsp* expression is sensitive to temperature. (A-D)** Head of MOC295 animals magnified at 20x using both bright field and epifluorescence microscopy. MOC295 animals carry the bipartite *hsp-16.41$_p$::cGAL/UAS$_p$::GFP* (green fluorescence). GFP fluorescence from *hsp-16.41* transcriptional activity can be detected in cells with axonal extensions (asterisks in C). MOC295 also carry the pan-neuronal transcriptional reporter *rab-3$_p$::2xNLS::TagRFP* (Stefanakis, Carrera, and Hobert 2015), visualised by red fluorescence (**D**). Arrowheads in B indicate overlap between green fluorescence from *16.41$_p$::cGAL/UAS$_p$::GFP* and red fluorescence in all neurons, indicating that under standard growth conditions neurons are the main tissue with a detectably active *hsp.41* promoter in somatic cells. Star in B indicates suspected GFP expression in the head mesodermal cell. (**E**) PS7171 young adult animals carrying *hsp-16.41*$_p$*::cGAL/UAS*$_p$*::GFP* were sorted into animals with either 0 to 1 GFP-expressing neurons or >3 GFP-expressing neurons. Levels of mRNA of GFP (coming from expression of transgene *hsp-16.41*$_p$*::cGAL/UAS*$_p$*::GFP*), as well as from endogenous *hsp-16.2* and *hsp-16.41* were measured by quantitative RT-PCR in high and low expressors. Paired t-test shows that there is a significant difference between the two groups of animals and that GFP expression from *hsp-16.41*$_p$*::cGAL/UAS*$_p$*::GFP* recapitulates endogenous *hsp-16.2* and *hsp-16.41* expression. **(F-G)** Images of PS7171 animals carrying bipartite expression system *16.41*$_p$*::cGAL/UAS*$_p$*::GFP* raised either at 15°C (F) or 25°C (G). Images were taken at 5X magnification. GFP expression in head and tail neurons is higher at 25°C than at 15°C. **(H-K)** Maximum intensity projection of 63x confocal images of the head region of *16.41*$_p$*::cGAL/UAS*$_p$*::GFP* worms treated for single-molecule RNA FISH (smFISH). The green channel shows hybridisation of a probe against *hsp-16* transcript. DAPI staining is in blue. (A,C) Worms grown at 15°C. (B,D) worms grown at 25° C (left panel). As shown above in (A-D), *hsp16.41* promoter is temperature sensitive. The smFISH experiment shows that animals with increased number of GFP-positive cells also have an increased number of endogenous *hsp-16* transcripts. **(L)** Quantification of smFISH shown in panel (G-J). Each dot represents number of head cells with *hsp-16* smFISH signal.
