## Supplementary material for "Inter individual variability in neuronal expression of heat shock protein genes predicts stress survival in *Caenorhabditis elegans*": Suplemental Methods

### SUPPLEMENTAL METHODS

#### 1-Nanofluidics-based qPCR for single worms improves sensitivity and accuracy of steady state mRNA detection.

##### 1.1-Parameter estimation and optimisation of the Worm-to-Ct qPCR method.

In qRT-PCR assays, gene expression is measured as threshold cycles ($Ct$). The latter are defined as the number of PCR amplification cycles that were required to achieve a given fluorescence level $F_0$ (Figure 1B, left). Often, qRT-PCR analyses, e.g., (Guo et al. 2010), adopt stepwise strategies in which $Ct$ values are pre-processed before downstream analyses (e.g. differential testing or clustering) are performed. After performing quality control to exclude poor quality samples, two pre-processing steps are typically applied: imputation of censored $Ct$ values and data normalisation. The latter refers to the removal of unwanted effects that can mask the signal of interest (Vallejos et al. 2017). These effects include intrinsic differences between samples (e.g. differences in mRNA content) and technical artefacts (e.g. uneven amplification). A popular strategy for qPCR assays is to impute censored data to then normalize it using the $\Delta Ct$ method, where one (or more) housekeeping (HK) genes are used as a reference and the geometric mean of a number of HK genes is used to normalise across samples. As discussed below, qPCR Bayes neither uses imputation for filtering nor the $\Delta Ct$ method for normalisation. Instead, it treats values that fall beyond the limit of detection (LOD) as undetermined and normalisation does not rely on the geometric mean of HK gene values, but instead on an integrative simultaneous approach. Despite these differences, qPCR Bayes requires a priori specification of LOD values and of HK control genes. This section details the methods used to estimate these parameters and also presents other essential qPCR controls that are required to subscribe to previously determined PCR guidelines (Bustin et al. 2009).

###### 1.1.1. Estimation of sensitivity and accuracy of Worm-to-Ct using RNA spike-in.

Estimating the limit of detection (LOD) is an essential step for defining the sensitivity of the assay. The LOD is defined as the lowest concentration at which 95% of the samples are detected (Bustin et al. 2009). Usually, LOD is determined by systematically reducing the number of PCR cycles until Cq/t values acquire a Poisson distribution. Instead of using this method, we determined the percentage of failed qPCR reactions as a function of the known concentration of added RNA spike-in. We used synthetic sequences that span a large linear detection, covering an 8292-fold range from 0.4578 to 3750 attomoles/µl, equivalent to a range that includes $10^6$ copies/ng of total RNA to rare transcripts of 120 copies/ng or less of total

RNA. The synthetic RNA spike-in used were established by the External RNA Control Consortium (ERCC) (Jiang et al. n.d.). The input concentration of spikes at which there is 5% dropout rate is considered the LOD, as previously described (Bustin et al. 2009). The failure rate increases exponentially at Ct values of 22 or higher (Fig. S1A) and therefore our estimation of LOD is 22 when using 10 amplification cycles.

To estimate the accuracy of the assay, it is necessary to establish the relationship between experimentally measured and actual concentrations. This can be performed using information provided by externally added synthetic RNA spike-in of known concentration. Figure S2B depicts the raw Ct values of a number of RNA spike-in amplified using the Worm to Ct protocol plotted against the number of RNA molecules added to the reaction. The graph shows a linear response across the entire detection range ($R^2$= 0.87, $p<0.0002$, F-test), demonstrating the high accuracy of the method. This linear regression is useful to estimate the range of detectable RNA concentrations, with the lower end corresponding to an input concentration of ~2 attomoles/μL of ERCC RNA spike-ins (equivalent to 100-200 copies/ng RNA) (Fig. S2B). The analysis shows that the qPCR run from the Worm-to-Ct method has a sensitivity that is comparable to what has been previously described (Devonshire, Elaswarapu, and Foy 2011) and can reliably detect Ct values superior or equal to 22, which is equivalent to 32 cycles on microliter-based PCR assay.

##### 1.1.2. Worm to Ct method provides cDNA, which is devoid of DNA contamination

Several steps must be considered in order to ensure the specificity of the assay, i.e. that the assay is detecting the appropriate target sequences. As described above, the primers used in this study have been tested for both specificity and efficiency. Yet another potential source of technical bias could be the presence of genomic DNA contamination in lysates obtained with the Worm-to-Ct protocol. Such contamination could lead to overestimation on the quantitation of steady state transcripts and therefore decrease the specificity of the assay. Although the number of copies of genomic DNA (gDNA) per gene in single worm samples (about 2000 copies) is significantly lower than in bulk samples, we directly estimated the efficiency of DNAse treatment. To assess potential DNA contamination on the samples, we run 2 arrays of a FlexSix IFC on a series of cDNA samples from 12 single worms versus minus reverse transcription (RT) controls from the same 12 worms. As shown in Figure S1C-F, no or negligible PCR amplification of DNA was obtained from single worm lysates lacking reverse transcription (-RT) for four different targets. This observation is true for primer sets that were designed to span exon-exon junctions (only present in cDNA obtained from mature mRNA transcripts) as well as for primer sets designed against exonic regions, which could amplify both genomic DNA (gDNA) and cDNA (all p-values<0.01, t-test). These results indicate that the DNAse treatment applied to Worm-to-Ct lysates is effectively eliminating gDNA contamination.

##### 1.1.3. Determination of the optimal set of genes used as normalisation factors.

The pipeline presented in this article does not rely solely on housekeeping (HK) genes for the normalisation of samples. However, it uses values of HK genes and therefore it is important to establish what the best HK genes are to use for the specific sample. We use young adult *C. elegans* in most of our assays and we therefore established the most stable HK genes to use at this specific stage using the geNorm algorithm (https://genorm.cmgg.be, (Vandesompele et al. 2002)), which identifies the most stable control genes for normalization (normalisation factors, NFs). This algorithm calculates the average pairwise variation of a particular NF with respect to the next $NF_{n+1}$ providing gene-stability measure “M”. Therefore, genes with the lowest M value have the most stable expression and higher ranking. We determined the M value for 7 potential NFs, commonly used in qRT-PCR assays, by running the geNorm algorithm on non-normalized expression data across experiments: 694 individual biological replicates across several genetic backgrounds and developmental stages (day 1 and day 2 of adulthood). Our geNorm analysis estimated a ranking of stability among potential normalising genes where *cdc-42* (M=0)> *pmp-3* (M=0.0320)> *ire-1*(M=0.0373)> *ama-1*(M=0.0517)> *ife-1*(M=0.0647)> Y45F10D.4(M=0.0750)> *gapdh/gpd-2*(M=0.0864). The geNorm algorithm also quantifies the optimal number of NFs, defined as the minimal number of control genes required to maintain the pairwise variation at its lowest (Vandesompele et al. 2002). Table 1 shows the optimal NFs is 3, as it already shows the lowest M, and using 3 or 7 NFs is almost equivalent. In our experiments we defined *cdc-42, pmp-3* and *ire-1* as the normalisation factors of choice. The primers sequences are listed in Table S1.

**Table 2: Determination of the optimal number of housekeeping genes using geNorm**

| **Number of remaining control genes** | **Mean M (stability value)** |
|---|---|
| 2 to 3 HK genes *(cdc-42+pmp-3 +/- ire-1)* | 0.012197 |
| 3 to 4 HK genes *(cdc-42+pmp-3+ire-1 +/-ama-1)* | 0.015629 |
| 4 to 5 HK genes *(cdc-42+pmp-3+ire-1+ama-1 +/-ife-1)* | 0.015524 |
| 5 to 6 HK genes *(cdc-42+pmp-3+ire-1+ama-1+ife-1 +/-Y45F10D.4)* | 0.014354 |
| 6 to 7 HK genes *(cdc-42+pmp-3+ire-1+ama-1+ife-1+Y45F10D.4 +/-gpd-2)* | 0.014930 |

##### 1.2-The single-worm Worm-to-Ct qPCR method provides better quality data than standard Trizol techniques.

In worm-to-Ct, worms are directly added to the lysis buffer, skipping the direct isolation of easily degradable RNA. To determine the quality of the data obtained by this method, we generated cDNA from 90 single worms. Half the worms were processed by the Worm-to-Ct method and the other half by the standard Trizol method. As highlighted in Figure S3A the percentage of Ct values per gene that are not censored using Worm-to-Ct method is higher than the percentage obtained using Trizol extraction. Likewise, the distribution of Ct values among non-censored observations tend to be on average lower when using Worm to Ct (Figure S3B). These observations indicate that the Worm to Ct method surpasses traditional Trizol extraction methods by increasing transcript capture efficiency from single worms and therefore decreasing the occurrence of false negatives.

#### 2- qPCR_Bayes: a Bayesian framework for differential variability analysis using qRT-PCR gene expression data.

##### 2.1-Statistical challenges when analysing qRT-PCR gene expression data

The output of qRT-PCR assays often includes *censored* observations in which $Ct$ values are “undetermined”. Censoring is particularly prominent when the input material for the assay is a small number of cells (Buettner et al. 2014) (e.g. single-worm or single-cell level). Censoring can be due to absence of expression, or to low gene expression. In practice, censoring arises in two scenarios: when the threshold $F_0$ is achieved but the observed $Ct$ value exceeds the limit of detection (LOD) after which $Ct$ values are deemed unreliable (Bustin et al. 2009) (Figure 1B, middle), or when the fluorescence threshold $F_0$ was not achieved (Figure S6B, right). A popular strategy to address censoring in the analysis of qRT-PCR data is to impute censored observations. Among others, such strategies include substituting censored $Ct$ values by a fixed quantity (e.g. LOD). However, it has been shown that imputation is sub-optimal in this context, as it can distort downstream analyses (Guo et al. 2010; Buettner et al. 2014; Pipelers et al. 2017).

A ubiquitous step when analysing data generated by gene expression assays (e.g. RNA sequencing, microarrays) is *normalisation* to eliminate unwanted effects such as cell size that can mask the signal of interest (Vallejos et al. 2017). A popular normalization strategy for qRT-PCR data is the $\Delta Ct$ method, where one (or more) housekeeping genes (HK) are used as a reference. For each gene $i$ and sample $j$, the $\Delta Ct$ method defines normalised values as $\Delta Ct_{ij} = Ct_{ij} - Ct_j^{HK}$, where $Ct_j^{HK}$ denotes the observed $Ct$ for the housekeeping gene in sample $j$. Alternatively, (Vandesompele et al. 2002) suggested to replace $Ct_j^{HK}$ by the geometric mean across multiple housekeeping genes. The $\Delta Ct$ method requires housekeeping genes to have stable expression across all samples. However, specifying such genes can be problematic as HK genes might vary across tissues or experimental conditions (Pérez-Novo et al. 2005). Another limitation of the $\Delta Ct$ method is that normalisation and other analyses (e.g. differential expression testing) are carried out sequentially – without propagating statistical uncertainty between these steps. The latter has been shown to damage the results of downstream analyses (Lewin et al. 2006; Wu and Irizarry 2007; Pipelers et al. 2017).

**2.2-qPCR_Bayes: an integrated approach to analyse qRT-PCR datasets.**

To address the limitations discussed above, we introduce qPCR_Bayes – an integrated Bayesian statistical framework which simultaneously: (i) addresses censoring using a probabilistic approach for which imputation of censored $Ct$ values is not required, (ii) performs built-in data normalisation which borrows information across all genes, and (ii) implements downstream analyses to robustly infer gene-level mean expression and expression variability within and between experimental conditions.

In Bayesian statistics, hyper-parameters, or parameter of a prior distribution before data is observed, need to be initialized. As shown in Figure 2A, we determined hyper-parameters by running synthetic data experiments with different hyper-parameters configuration and comparing to values obtained from an experimental data set for both mean and variability (see more detailed explanation in prior specification in materials and methods).

The layout of nanofluidics chips for high-throughput qRT-PCR allows monitoring of gene expression in a maximum of 96 single worms samples. To determine the minimum reasonable sample size to allow accurate monitoring of inter-individual gene expression variability, we used *in silico* simulations of experimental data obtained for 90 single worms, spanning different sub-sample sizes (Figure 2B-E). Based on this, we estimated a sample size of n=45 single worms to be reasonable, with a spearman correlation coefficient between simulated and experimental data >0.9995 for mean and >0.986 for variability (Figure 2C). This conveniently allows testing two conditions simultaneously on the same nanofluidics chip. Figure 2D-E shows good concordance of mean and variability measured for two groups of simulated conditions each including n=45 random samples among the pool of 90 worms.

In qPCR_Bayes, multiple *control* genes are used to guide the normalisation and quantify background technical noise. Firstly, in line with the $\Delta Ct$ method, housekeeping genes are used as an internal control to capture intrinsic differences in mRNA content and technical variability (e.g. amplification biases) across samples. When extrinsic RNA spike-in (e.g. the ones introduced in (Jiang et al. n.d.) are included in the assay, qPCR_Bayes uses this information to disentangle intrinsic from technical differences across samples (Vallejos et al. 2017). Throughout, *target* genes refer to all remaining non-control genes. Thus, one of the advantage of qPCR_Bayes is that it can use two different methods, $\Delta Ct$ method or imputation method, to normalise for sample specific effects. Figure S3 shows a comparison of qPCR_Bayes using either HK genes as normalisation factors, or using sample-specific parameters as normalisation factors to account for global differences between samples (implementation method). The integrated strategy for normalisation has advantages over the ΔCt method, particularly for samples with unusual behaviour of HK genes, as illustrated for the problematic effect of ΔCt method on outliers in Figures 3C and D. The data is overall more robust, with higher Pearson's (ρ) and Spearman's (r) correlations with respect to the estimates obtained when using all three genes as housekeeping genes (Figure 3E).

In qPCR_Bayes, the presence of censoring information in the data allows to use a probalistic framework inspired by survival analysis models and this removes the need for pre-processing imputation step (Pipelers et al. 2017; Buettner et al. 2014). The likelihood contribution of censored values is set, using the LOD as an experimental limit of detection, as the probability of observing a Ct value that exceeds the LOD.

Let $Ct_{ij}$ be the observed $Ct$ value for gene $i$ in sample $j$. If spike-in genes are available, the statistical model implemented in qPCR_Bayes is defined by:

$$Ct_{ij} = \begin{cases} \phi_j + s_j + \mu_i + \rho_{ij} + \epsilon_{ij}, & \text{if gene } i \text{ is target,} \\ \phi_j + s_j + \mu_i + \epsilon_{ij}, & \text{if gene } i \text{ is housekeeping ,} \\ s_j + \mu_i + \epsilon_{ij}, & \text{if gene } i \text{ is a spike-in,} \end{cases}$$

**Eq. 1**

where

$$\rho_{ij} \mid \delta_i \sim \mathrm{N}(0, \delta_i) \quad \text{and} \quad \epsilon_{ij} \mid \sigma^2 \sim \mathrm{N}(0, \sigma^2).$$

**Eq. 2**

If spike-ins are not available, a simplified model is defined by replacing Equation 1 with

$$Ct_{ij} = \begin{cases} \phi_j + \mu_i + \rho_{ij} + \epsilon_{ij}, & \text{if gene } i \text{ is target,} \\ \phi_j + \mu_i + \epsilon_{ij}, & \text{if gene } i \text{ is housekeeping.} \end{cases}$$

**Eq. 3**

A graphical representation of these models is displayed in Figure 1C.

If censoring is present in the data, the models above are extended using a probabilistic framework that is motivated by survival analysis models. Formally, observed $Ct$ values are denoted as $Ct_{ij} = \min\left(Ct^*_{ij}, LOD\right)$, where $Ct^*_{ij}$ is a latent unobserved value and LOD is an experimental limit of detection (Lewin et al. 2006; Bustin et al. 2009). As such, the likelihood contribution of censored values is set as the probability of observing a $Ct$ value that exceeds the LOD. As shown in (Pipelers et al. 2017; Buettner et al. 2014), this removes the need for a pre-processing imputation step.

In qPCR_Bayes, sample-specific parameters act as normalisation factors that capture global differences between samples. To infer these parameters, qPCR_Bayes borrows information across all target and control genes. In Equation 1, the use of synthetic ERCC RNA spike genes enables the qPCR_Bayes to disentangle two types of effects: while $s_j$ captures technical effects that affect all genes (e.g. amplification biases), $\phi_j$ captures intrinsic differences in mRNA content that effect only housekeeping and target genes. The latter can be due, for example, to global whole-transcriptome upregulation (Lovén et al. 2012) or differences across cell-cycle stages (Buettner et al. 2015). Instead, in Equation 2, a single set of normalization parameters $\phi_j$ captures a combination of intrinsic and technical sample-specific effects

In Equations 1 and 2, gene-specific parameters $\eta_i = 2^{-\mu_i}$ can be interpreted in terms of mean expression levels for the population under study ($Ct$ values are in $\log_2$ scale). Moreover, the residual term $\epsilon_{ij}$ captures background technical noise that is shared by all genes and samples. The strength of this variability is controlled by a global variance parameter $\sigma^2$. Finally, the random effects $\rho_{ij}$ capture biological sample-to-sample variability of target genes, whose strength is controlled a gene-specific parameter $\delta_i$. We observed that both models (with and without spike-in genes) led to comparable posterior inference for gene-specific parameters (data not shown).

As in (Vallejos, Richardson, and Marioni 2016), qPCR_Bayes is extended to identify differences in gene expression patterns between pre-specified experimental conditions (groups). For this purpose, we assume that gene-specific parameters are also group-specific.

For each group $p$, we denote these as $\eta_i^{(p)}$and $\delta_i^{(p)}$. When comparing two groups $p$ and $p'$, changes in expression variability are then quantified as the $\log_2$ fold change between group-specific variance parameters, $\log_2\left(\delta_i^{(p)}/\,\delta_i^{(p')}\right)$. Similarly, mean expression differences between the groups are quantified as $\log_2\left(\eta_i^{(p)}/\,\eta_i^{(p')}\right) = \mu_i^{(p')} - \mu_i^{(p)}$. In both cases, statistically significant changes are highlighted using a probabilistic decision rule that is calibrated by controlling the expected false discovery rate (Roberts and Rosenthal 2009).

Recently, (Pipelers et al. 2017), introduced a statistical method to perform differential (mean) expression tests that uses a similar approach to deal with censored observations and to carry out data normalisation. However, the differential test implemented in qPCR_Bayes is more general, also enabling differential variability analyses. Moreover, our implementation uses a Bayesian framework to generate a joint posterior distribution for all model parameters (see implementation details below). As such, our differential tests do not rely on asymptotic properties of maximum likelihood estimators.

#### 2.3-Implementation details

##### 2.3.1- Identifiability

The models introduced by Eq. 1 and Eq. 3 are not identifiable In other words, for any arbitrary constant $\tau$, the data does not contain enough information to distinguish between parameter values $(\phi_j,\ \mu_i)$ and $(\phi_j+\tau,\ \mu_i-\tau)$. As a solution, we impose an identifiability constraint: we assume that $\mu_i$ is known for all housekeeping and spike-in genes. While these values are not known a priori, in practice, we set these to be equal to the average $Ct$ value that is observed for each gene (excluding censored values).

##### 2.3.2- Prior specification and parameter estimation

Independent prior distributions are assigned to all model parameters. For each target gene, the prior distribution of gene-specific parameters $\mu_i$ is set as

$$\mu_i \sim \mathrm{N}\left(m_0, s_\mu^2\right) \quad \text{and} \quad \delta_i \sim \text{Student } t_\nu\left(0, s_\delta^2\right).$$

In the spirit of empirical Bayes, $m_0$ is set as the average observed $\mathrm{Ct}$ value across all target genes. Based on synthetic data experiments (Figure 2A), default values for the remaining hyper-parameters were set as $s_\mu^2 = s_\delta^2 = 2$ and $\nu = 5$. As shown in Figure 2, these values shown good empirical performance across different sample sizes.

The following prior distributions are assigned to the remaining model parameters

$$\phi_j \sim N(0, s_\phi^2), \quad s_j \sim N(0, s_s^2) \quad \text{and} \quad \sigma^2 \sim \text{Inv-Gamma}(a_{\sigma^2}, b_{\sigma^2}).$$

In these prior distributions, we arbitrarily set $s_\phi^2 = s_s^2 = a_{\sigma^2} = b_{\sigma^2} = 1$ as a default choice. However, the posterior distribution was stable upon changes in these values.

The posterior distribution for all model parameters was obtained using an adaptive Metropolis within Gibbs algorithm (Roberts and Rosenthal 2009). Our implementation is freely available as an R package (R Core Team 2018). This package can be found at https://github.com/catavallejos/qPCRBayes.

##### 2.3.3- Post-hoc offset correction between groups

The role of the sample-specific parameters $\phi_j$ is to capture intrinsic sample-to-sample differences (e.g. mRNA content) within a single experimental condition. If the goal of the analysis is to perform differential expression analyses between two experimental conditions, it is also important to adjust for global changes in expression (e.g. global changes in mRNA content) between them. As in (Vallejos, Richardson, and Marioni 2016), between groups normalisation is implemented through a post-hoc offset correction. This strategy assumes that the average expression level of housekeeping genes is the same for all groups of samples. Let $Ct_{ij}^{(p)}$ be the observed $Ct$ value for housekeeping gene $i$ in sample $j$ for group $p$, $n_p$ be the number of samples in group $p$ and $N_{HK}$ be the total number of housekeeping genes. Moreover, let $\mu_i^{(p)}$ denote gene-specific mean parameters within group $p$ and $\phi_j^{(p)}$ denote sample-specific normalisation parameters within group $p$. A post-hoc offset correction for these parameters is defined as follows:

$$\tilde{\mu}_i^{(p)} = \mu_i^{(p)} - \Lambda_p \text{ and } \tilde{\phi}_j^{(p)} = \phi_j^{(p)} + \Lambda_p,$$

where $\Lambda_p = \frac{1}{N_{NK}} \sum_{\text{i is housekeeping}} \left[\frac{1}{n_p} \sum_{j=1}^{n_p} C_{ij}^{(p)}\right]$.

For simplicity, censored $Ct$ values are excluded from this calculation. This decision is not problematic as samples for which the expression of housekeeping genes is not captured (censored) are typically removed during the quality control step.

For ease the notation, once this offset correction has been applied, we use $\mu_i^{(p)}$ and $\phi_j^{(p)}$ to denote $\tilde{\mu}_i^{(p)}$ and $\tilde{\phi}_j^{(p)}$, respectively.

##### 2.3.4- Differential expression testing

Statistical evidence for changes in gene expression is quantified through tail posterior probabilities associated to large log2 fold changes in mean and/or variability between experimental conditions. Let $\mu_i^{(p)}$ and $\mu_i^{(p')}$ represent mean $\mathrm{Ct}$ parameters for two groups of samples. For a given posterior probability threshold $\alpha_M$ $(0.5 < \alpha_M < 1)$, a gene $i$ is identified to be differentially expressed if

$$\pi_i^M(\tau_0) = \mathrm{P}\left(\left|\mu_i^{p'} - \mu_i^{p}\right| > \tau_0 \,\middle|\{\text{data}\}\right) > \alpha_M,$$

where $\tau_0$ corresponds to a minimum tolerance log2 fold change threshold. Similarly, for a given posterior probability threshold $\alpha_V$ $(0.5 < \alpha_V < 1)$, a gene $i$ is identified to be differentially variable if

$$\pi_i^V(\omega_0) = \mathrm{P}\left(\left|\log_2\left(\delta_i^{(p)}/\,\delta_i^{(p')}\right)\right| > \omega_0 \,\middle|\{\text{data}\}\right) > \alpha_V,$$

where $\omega_0$ is a minimum tolerance log2 fold change threshold. In the limiting case, where $\tau_0 = 0$ or $\omega_0 = 0$, the decision rules above are modified as in (Bochkina and Richardson 2007).

Posterior probability thresholds $\alpha_M$ and $\alpha_V$ are calibrated to achieve a desired expected false positive rate (EFDR, (Newton, Noueiry, and Sarkar 2004)). Values for $\tau_0$ and $\omega_0$ must be set a priori. As a default, we set $\tau_0 = \log_2(1.5)$ and $\omega_0 = \log_2(2) = 1$. For the sample sizes adopted in our experiments, these values showed good empirical performance – i.e. they covered the range of log2-fold changes that can be expected by chance.

Finally, to avoid possible confounding effects between mean and variability estimates, our default option is to exclude those genes identified to exhibit a change in mean expression when assessing changes in variability.
